# Cancer/Testis genes are predictive of breast tumor subtypes

**DOI:** 10.1101/2021.10.27.465656

**Authors:** Marthe Laisné, Sarah Benlamara, André Nicolas, Lounes Djerroudi, Nikhil Gupta, Diana Daher, Laure Ferry, Olivier Kirsh, Claude Philippe, Yuki Okada, Gael Cristofari, Didier Meseure, Anne Vincent-Salomon, Christophe Ginestier, Pierre-Antoine Defossez

**Affiliations:** EDC; Institut Curie; IRCAN, Nice; Tokyo University; CRCM

## Abstract

Breast cancer is the most prevalent type of cancer in women worldwide. Within breast tumors, the basal-like subtype has the worst prognosis and no dedicated therapy, therefore new tools to understand, detect, and treat these tumors are needed. Certain germline genes are re-expressed in tumors, and constitute the Cancer/Testis genes; their misexpression has diagnostic and therapeutic applications. Here, we designed a new approach to examine Cancer/ Testis gene misexpression in breast tumors. We identify several new markers in Luminal and HER-2 positive tumors, some of which predict response to chemotherapy. We then use machine learning to identify the 2 Cancer/Testis genes most associated with basal-like breast tumors: HORMAD1 and CT83. We show that these genes are expressed by tumor cells but not the microenvironment, and that they are not expressed by normal breast progenitors, in other words their activation occurs de novo. We find these genes are epigenetically repressed by DNA methylation, and that their activation upon DNA demethylation is irreversible, providing a memory of past epigenetic disturbances. Basal-like tumors expressing both genes have a poorer outcome than tumors expressing either gene alone or neither gene. Therefore, these findings suggest a potential synergistic effect between Cancer/Testis genes in basal breast tumors; these findings have consequences for the understanding, diagnosis, and therapy of the breast tumors with the worse outcomes.

## INTRODUCTION

Cancer cells undergo massive genetic and epigenetic changes relative to their normal progenitors. The advances of genomics and epigenomics have yielded an ever more complete picture of these abnormalities, and drawn accurate molecular portraits of different tumor types. The large number of samples examined in public cohorts increase statistical power, yet parsing out driver from passenger events remains far from trivial (Muiños F. et al., 2021).

Altered gene expression is one of the functional consequence of genetic and epigenetic modifications in tumors. Genes can be turned off by deletions, alterations in their control elements such as enhancers, or changes in the transcriptional machinery. Conversely, they can become overexpressed by amplification, gain of enhancers, or expression of transcriptional activators, among other possibilities. Genes that are frequently turned on in a tumor type are useful as biomarkers. In some instances, their expression can inform prognosis and choice of treatment. Finally, these over-expressed genes can play a physiological role in the tumor cells, and therefore represent therapeutic targets. HER2 is such an example: the gene can be amplified, its overexpression marks a specific subtype of breast tumors, and highly efficient therapeutic antibodies have been generated against this target.

HER2 is expressed by normal breast cells, so its overexpression in breast tumors is just the amplification of a pre-existing expression pattern. However, tumor cells can also deviate radically from their ancestral gene expression pattern and turn on genes that are normally activated in other tissue types or other developmental stages (Wang J. et al. 2014). For instance, various tumor types, in men and women, express genes that are typical of the placenta (Rousseaux S. et al. 2014; Naciri et al. 2019). Within this broad framework of ectopic gene reactivation in tumors, one class of genes bears special conceptual interest and therapeutic promise: the cancer/testis genes.

As their name implies, the cancer/testis genes are normally expressed only in the male germline, but become reactivated in tumors, both in female and male patients (Whitehurst AW 2014). As they are not expressed in any normal somatic cells, they are remarkable biomarkers for tumors. In addition, as the testis is an immune sanctuary in men, and as the testicular genes are not normally expressed in women, their expression in tumors opens an excellent possibility for immunotherapy. Finally, cancer/testis genes may be oncogenes in their own right, and are potential drug targets for therapy (Gibbs ZA & Whitehurst AW 2018).

Breast cancer is the most common cancer in women, both in developed and developing countries, and breast malignancies killed almost 700,000 women worldwide in 2020 (www.who.int). It has long been appreciated that breast tumors form an heterogeneous ensemble, with at least 5 distinguishable subtypes: normal-like, Luminal A, Luminal B, HER2-positive, and basal-like. Within those groups, basal-like tumors could themselves contain distinct subtypes, and they have the worst prognosis and no dedicated therapy.

Cancer/testis genes have been investigated as potential biomarkers, oncogenes, and targets in breast cancer, with promising results (Kaufmann J. et al. 2019; Paret C. et al. 2018; Adams S. et al. 2011; Mischo A. et al. 2006). To build on these investigations, we undertook an unbiased analysis of publically available expression data with a new bioinformatic approach. This led us to discover several new markers associated with different breast tumor subtypes. Our cohort of in situ tumors establishes that cancer/testis gene activation is an early event in tumorigenesis, and that there is no switch of their expression pattern between early and more established tumors. We then focused on the two genes whose expression is most highly associated with basal breast tumors: HORMAD1 and CT83. We show that these genes are not expressed by healthy progenitors, but expressed de novo in the tumor cells. We demonstrate that loss of methylation is sufficient to reactivate both genes, and that an initial activation event is sufficient to trigger persistent expression. Most basal tumors express at least one of the two genes, but those that express both have significantly worse outcome, hinting at a cooperative effect. These findings advance our conceptual understanding of cancer/testis genes in breast cancer, and they have practical implications for diagnosis and treatment. These results also suggest new experiments to understand the potential synergistic effect of HORMAD1 and CT83 co-activation in breast cancer tumorigenesis.

## RESULTS

### A custom bioinformatic approach identifies the Cancer/Testis genes most associated with breast tumors

The first step of our study was to establish an exhaustive list of C/T genes; it includes all of the C/T genes described in three independent publications, for a total of 1350 genes (Almeida et al. 2009; Rousseaux et al. 2013; Wang et al. 2016). Our second resource was genomics data, including RNA-seq, from The Cancer Genome Atlas (TCGA), covering 1090 tumors samples and 113 healthy juxtatumoral mammary samples (Figure 1A).

**Figure 1:**
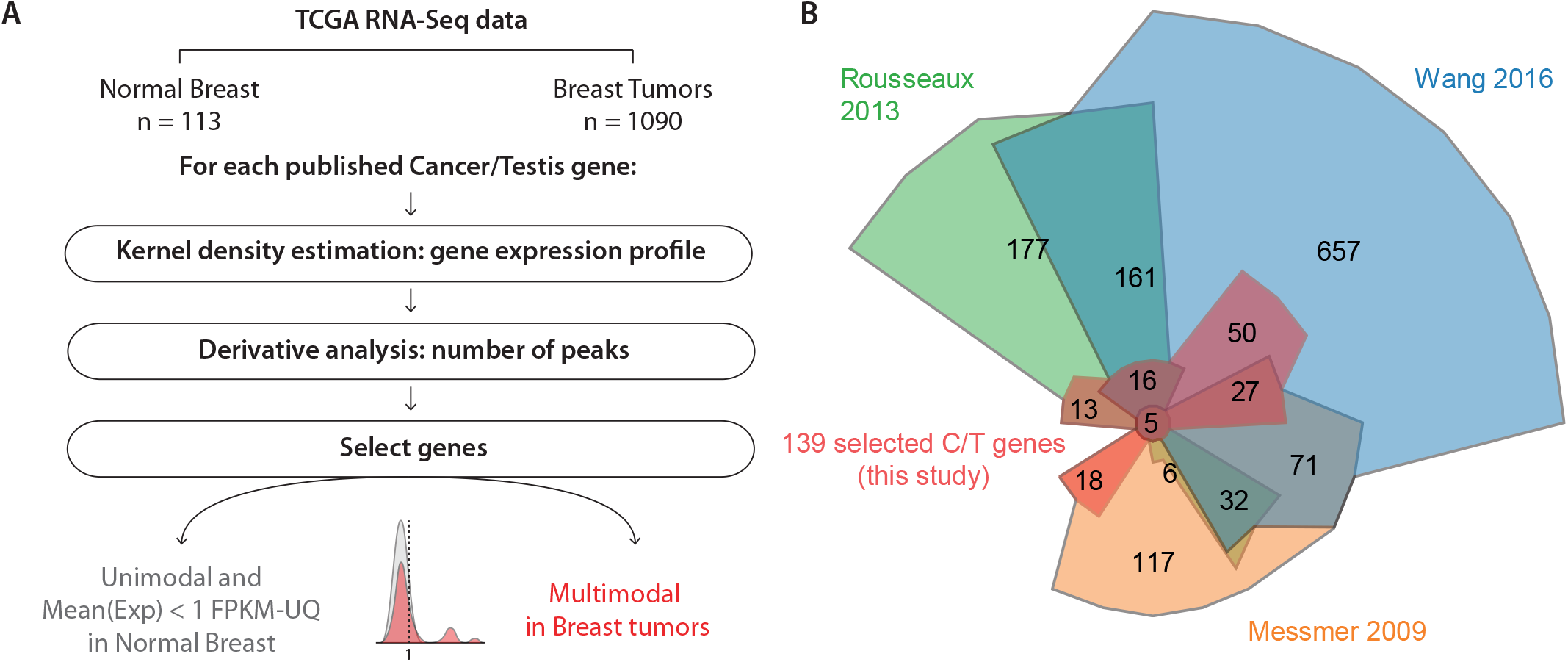
A custom bioinformatic screen identifies 139 Cancer/Testis genes abnormally expressed in breast tumors. A. Schematic description of the bioinformatic pipeline. We depict the expression profile of a gene that passed the screen: it has a unimodal, zero-centered profile in normal tissue, and a multimodal profile in breast tumors. B. Chow-Ruskey diagram showing the intersection between previously published C/T gene lists and the C/T genes that were selected for our study.

To identify C/T genes reactivated in breast tumors, we established a custom bioinformatic approach. An ideal biomarker should have little or no expression in healthy samples, but high expression in at least some of the tumors. Mathematically, these properties are reflected in a zero-centred, single-mode density function in healthy breast samples, and a multi-mode density function with one or more non-zero maxima in tumor samples, reflecting one or more groups of tumors that have activated this gene. Such profiles can be detected automatically by examining changes in the derivative of the density function (Figure 1A).

To implement this idea, we created a two-step pipeline. First, we determined the distribution of expression of each C/T gene in healthy mammary samples and in breast tumors, and smoothed these distributions using kernel density estimation. As it is crucial to not overfit or oversmooth expression values, we systematically tested multiple values for the bandwidth parameter using positive and negative controls (Figure S1A) and we selected a balanced value (band-width = 0.7). Second, we analyzed the derivative of the distribution function to obtain the number of distinct peaks. This allowed us to focus on C/T genes that are not expressed in healthy mammary samples (unimodal expression profile centered on 0 according to kernel density estimation), but activated in some breast tumor samples (multimodal expression profile).

Our method complements previously used approaches in that it is orthogonal, less calculation-intensive, flexible, and sensitive. Of note, this unbiased scheme is not restricted to C/T genes and it could be broadly used to identify any other genes that are abnormally expressed in tumor samples compared to matched normal juxta-tumor tissues, such as potential tumor suppressor genes or oncogenes (Figure S1B-D). Our approach allowed us to define a highly selective list of 139 C/T genes with abnormal expression profile in breast tumors compared to normal breast (Figure 1B, Supplementary Table 1). The examination of GTEx RNA-seq data confirmed that these 139 genes are expressed in the human germline, but not in the breast (or other healthy tissues, Figure S1G). The reactivation seen in tumors is therefore a pathological event.

### The activation of selected C/T genes marks different subtypes of tumors and cell lines

We then tested whether the expression of certain members of our 139-gene list was specifically associated with certain subtypes of breast tumors. For this, we used Principal Component Analysis (PCA) on TCGA data, using the subtype annotations provided for each tumor (Figure 2A). A visual inspection suggested that tumor types could be separated on the basis of C/T gene expression (Figure 2A), with a clearly distinct group of basal tumors, for instance. These clusters were also found when the tumors were classified on the basis of their anatomohistological subtype, rather than their transcriptome-defined subtype (Figure S2A), and they were also visible when UMAP was used instead of PCA (Figure 2A, S2A). We therefore conclude that expression of some genes in our list can stratify breast tumors by subtypes.

**Figure 2:**
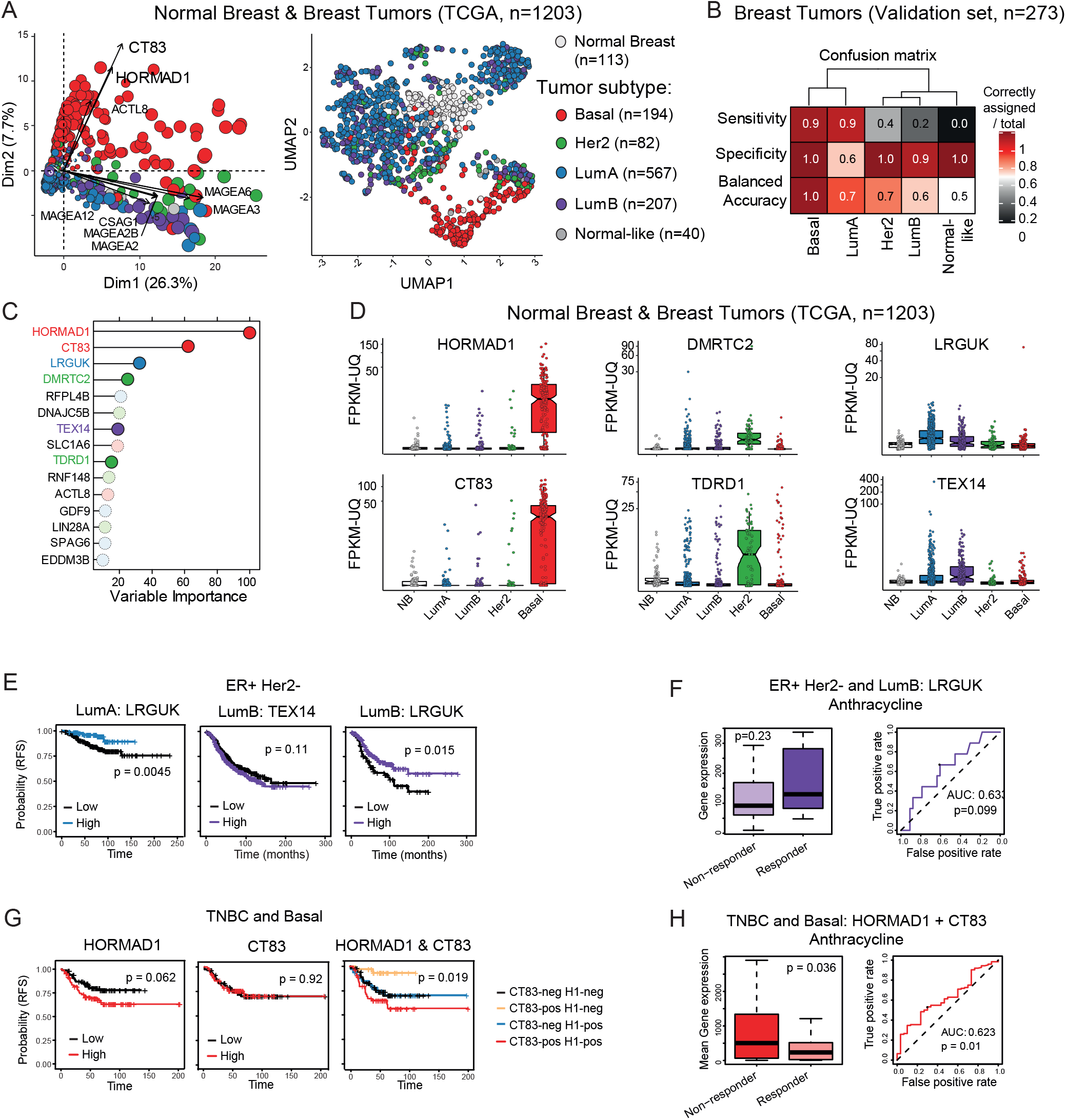
The activation of specific C/T genes is predictive of tumor subtype, occurs early during tumorigenesis, and is associated with prognosis. A. Multidimensional analysis of TCGA breast tumor and healthy samples based on expression of the 139 selected C/T genes. Each dot is a sample, the color code corresponds to the tumor subtype by PAM50 molecular classification. Left: Principal Component Analysis, dot sizes are proportional to the quality of representation in PC1/ PC2 space. The C/T genes best correlated to PC1/PC2 are represented. Right: Uniform Manifold Approximation and Projection (UMAP). B. Confusion matrix for breast tumor samples in the validation cohort (25% of the samples, randomly selected from the TCGA breast tumors), using the the best Random Forest model. This model was established after a 500-tree training on the discovery cohort (75%), based on the expression level of the 139 C/T genes. C. The top 15 most important variables in the best Random Forest model for PAM50 subtype prediction. The color of the gene name indicates the tumor type most associated. D. Expression levels for 6 subtype-specific C/T genes in the breast TCGA cohort, according to PAM50 tumor sub-type. E. Relapse-free survival curves for ER+ Her2-breast cancer patients according to LRGUK expression, for Luminal A tumors (left), and for Luminal B tumors (right). F. Left: Expression value for the luminal-specific C/T gene LRGUK in luminal B tumors, according to the clinical evaluation of tumor response to chemotherapy. Right: ROC curve evaluating the potential of LRGUK as a predictive biomarker of anthracyclin chemotherapy response of ER+ Her2- Luminal B tumors. G. Relapse-free survival curve for ER-PR-Her2- Basal-like breast cancer patients, as a function of HORMAD1 expression alone, CT83 expression alone, or combined expression of the two C/T genes. H. Left: Combined expression value for the two basal-specific C/T genes HORMAD1 and CT83 in basal-like tumors, according to the clinical evaluation of tumor response to chemotherapy. Right: ROC curve evaluating the potential of HORMAD1 and CT83 combined expression as a predictive biomarker of anthracyclin chemotherapy response of ER- PR- Her2- Basal-like tumors.

To identify these genes systematically we used a machine learning approach. We established a random forest model on a training set of TCGA breast tumors (75% of all samples, n=817), and tested the best model on the remaining tumors (n=273). This model could very effectively identify basal tumors, with high sensitivity (0.9) and high specificity (1.0), leading to a balanced accuracy nearing 100% (2B). Again, similar results were found when the tumors were classified anatomopathologically, rather than transcriptionally (Figure S2B). For Luminal B and Her2 subtypes the specificity scores were high (1.0 and 0.9 respectively), but the sensitivity lower (0.4 and 0.2) (Figure 2B). This could be due to the fact that some tumors of these groups do not express any C/T genes, leading to a lack of available information for the prediction.

Using the best random forest model, we ranked the 139 C/T genes according to their predictive value; the top 15 C/T genes are depicted in Figure 2C (and in Figure S2C for the analysis carried out with anatomopathological stratification). The two best predictors, HORMAD1 and CT83, are strongly associated with basal breast tumors: of the 190 basal-like breast tumors, 89% expressed either HORMAD1 or CT83, compared to only 13% of HER2-amplified, 6% of Luminal B, and 2% of Luminal A tumors (Figures 2D and S2D). These results are consistent with several previous reports that have associated HORMAD1 or CT83 expression with basal tumors (Watkins et al. 2015; José Adélaïde et al. 2007; Chen et al. 2019; Paret et al. 2015; Kondo et al. 2018; Chen et al. 2021), and they validate our approach. HORMAD1, a gene on human chromosome 1q21.3, is physiologically expressed by the pre-leptotene spermatocytes (Shin et al., 2010) and it regulates meiotic progression. CT83, on the other hand, is located on human chromosome region Xq23, it is expressed in mature sperm (Jung et al., 2019) but its reproductive function is unknown.

The expression of two other markers, DMRTC2 and TDRD1, is associated with HER2-positive tumors (Figure 2D), but the association is looser than that of HORMAD1/CT83 with basal tumors. During spermatogenesis, DMRTC2 has essential functions during pachytene (Date et al. 2012), whereas TDRD1 interacts with piRNAs and Piwi proteins to promote silencing (Mathioudakis et al. 2012). To the best of our knowledge, neither DMRTC2 nor TDRD1 have been previously linked to breast cancers in general, and to the HER-2 positive subtype in particular.

Lastly, we found two markers, LRGUK and TEX14, for which expression tends to mark Luminal tumors (Figure 2D). LRGUK is involved in diverse aspects of sperm assembly, including the microtubule-based shaping of spermatozoids (Liu et al. 2015); it was more frequently over-expressed in luminal A breast tumors (Figure 2D). As for TEX14, a factor necessary for intracellular bridges in germ cells (Greenbaum et al. 2006), it marked luminal B breast cancers, as well as luminal A tumors to a smaller extent (Figure 2D). While TEX14 has previously been linked to basal breast tumors (Karlin et al. 2015), we believe we present the first report that is actually much more prevalently expressed in Luminal tumors, especially of the more aggressive B subtype, and we are not aware of any publications linking LRGUK to breast tumors in general, nor to Luminal tumors in particular.

We next tested whether the associations we had detected using tumor expression data also held true with cancer cell lines. For this, we determined the expression level of the 6 markers described above in all the breast cell lines found in the Cancer Cell Line Encyclopedia (Figure S2E). We observed a good general agreement between tumors and cell lines of the same subtype. For instance, HORMAD1 and/or CT83 were highly expressed in the basal cell lines such as MDA-MB-436, MDA-MB-468, and HCC1599, but not in Luminal or HER2-positive cells. DMRTC2 and/or TDRD1 expression marked HER2-positive lines like AU565 or SKBR3. Finally, a typical Luminal A line, MCF7, expressed LRGUK and TEX14.

### Marker expression can be associated with response and survival

Finally, we asked whether the expression of these CT genes could distinguish, within a breast cancer subtype, tumors with a different prognosis or therapeutic response. We examined relapse-free survival at more than 10 years, on a large panel of breast tumors of known subtype (Győrffy 2021).

Activation of LRGUK in Luminal A or Luminal B tumors, was an indicator of good prognosis (Figure 2F). Furthermore, activation of the gene tended to correlate with better response to anthracyclines, although the trend failed to reach significance (Figure 2E).

For Her2-positive tumors, the expression of TDRD1 was not statistically linked to survival, whereas DMRTC2 expression correlated with poorer survival (Figure S2F). To detect other potentially useful characteristics of these tumors, we examined their immunological signature with the Immunoscore tool (Bindea et al. 2013) (Figure S2G): those with high DMRTC2 were expected to be more “hot”, i.e. more infiltrated, but also could be more immunosuppressive (high FOXP3 activation). Therefore, they might be attractive candidates for treatment with immune checkpoint inhibitors (Galon et al. 2019). As far as we are aware, all of these associations are new and may be helpful for prognosis and treatment choice.

The situation was particularly interesting for HORMAD1 and CT83 in basal-like tumors (Figure 2G). Neither gene considered alone was associated with prognosis, however the co-expression of both genes led to a significantly worse outcome, hinting at a possible synergistic effect. In addition, expression of both genes simultaneously correlated with a poorer response to anthracycline chemotherapy (Figure 2H).

### HORMAD1 and CT83 mark are expressed by most cancer cells in basal-like tumors, but are not expressed by the microenvironment

As basal-like tumors are especially deadly, we aimed the rest of our investigations on this tumor type. We started by repeating our random forest analysis on RNA-seq data from an independent set of tumors (Varley et al. 2014). In that second cohort also, HORMAD1 and CT83 were the most informative genes, and the most associated with basal tumors (Figures 3A and 3B). This independent cohort further supports the relevance of these 2 genes in basal tumors, thus we focused on HORMAD1 and CT83 in the rest of our work.

**Figure 3:**
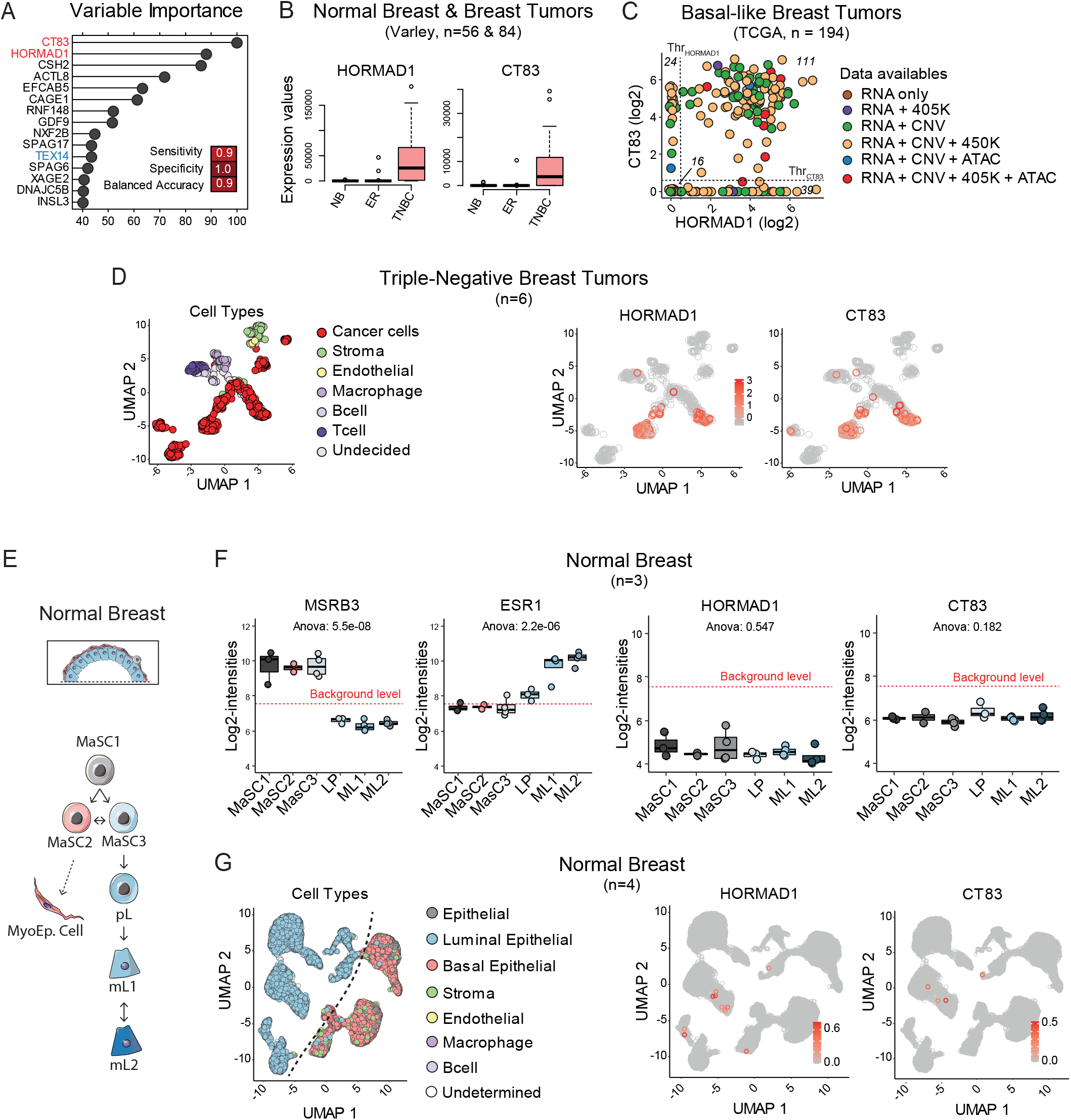
HORMAD1 and CT83 are expressed specifically by cancer cells, however scRNA-seq reveals rare HORMAD1+ / CT83+ luminal progenitor cells in healthy mammary gland. A. Top 15 most important variables in the best Random Forest model applied to an independent cohort of breast tumors. B. Expression of HORMAD1 and CT83 in the indicated sample types of the Varley/Myers cohort (GSE58135) C. Co-expression of *HORMAD1* and *CT83* based on RNA-seq analysis (log2 FPKM-UQ) in basal-like breast tumor samples (n=194) from the TCGA. Threshold for positive or negative expression are calculated based on the corresponding gene expression profile in tumors at the second inflexion point of the representative curve. The number of tumors belonging to each category is shown. D. UMAP representation of a scRNA-seq study on 6 triple-negative breast tumors (GSE75688). Each dot is either a tumor cell or a cell from the tumor microenvironment. From left to right: cell types which were determined based on the expression of specific marker genes; *HORMAD1* normalized expression level; *CT83* normalized expression level. E. Schematic representation of the mammary cell hierarchy in healthy adult mammary gland. *F. HORMAD1* and *CT83* expression in sorted healthy mammary cells. The red dotted line represents the threshold for gene expression detection. G. UMAP representation of a scRNA-seq study on 4 healthy mammary glands (GSE113197), after an enrichment in epithelial cell by FACS. From left to right: cell types which were determined based on the expression of specific marker genes; *HORMAD1* normalized expression level; *CT83* normalized expression level.

In the TCGA cohort, ∼90% of basal-like tumors expressed HORMAD1 or CT83 at the RNA level, and ∼60% expressed both (Figure 3C). Basal-like tumors are a heterogeneous ensemble, but tumors expressing both HORMAD1 and CT83 tended to form a more homogeneous set, with fewer distinct anatomopathological groups and a reduced number of molecular signatures (Figure S3A, Supplementary Table 2). Using the Lehmann classification (Lehmann et al. 2016), we found double-positive tumors in all subgroups except for Luminal Androgen Receptor (Figure S3B). In breast cancer cell lines as well, 70% of basal-like cell lines from CCLE were positive for HORMAD1 and/or CT83 (Figure S3C).

To verify that tumor cells themselves expressed HORMAD1 and CT83 (and not non-tumor cells of the microenvironment), we re-analyzed previously published single-cell RNA-seq data of 6 triple-negative breast tumors (of which 5 express HORMAD1 and CT83) (GSE75688, Chung et al. 2017). We found very clearly that only tumor cells (and not the microenvironment) express HORMAD1 and/or CT83 (Figure 3G). Within any given tumor, approximatively 20-40% of individual cancer cells express either HORMAD1 or CT83, and around 5-20% express both ; but we have to keep in mind that approximatively 50% of mRNA molecules are lost by scRNA-seq. Immunohistochemistry analysis of tumor samples from breast cancer patients will be more accurate for this point.

Taken together, these results at the RNA and protein level show that HORMAD1 and CT83 are expressed by most tumoral cells in most basal-like tumors, and that they are not expressed by the microenvironment.

### Most healthy mammary cells fail to express HORMAD1 or CT83

As HORMAD1 and CT83 are expressed by tumor cells, and as these tumor cells derive from the transformation of healthy breast cells, we asked whether the 2 genes are expressed by progenitors found in healthy breast. For this, we turned to RNA expression data obtained on healthy cells sorted from reduction mammoplasties, where markers were used to FACS-sort stem cells, luminal progenitors, and mature luminal cells (Figure 3E, Morel et al. 2017). Known genes displayed the expected expression pattern: for example MSRB3 was expressed in stem but not more differentiated cells, whereas ESR1 had the opposite pattern (Figure 3F). In contrast, neither HORMAD1 nor CT83 was detectably expressed in any of the sorted cell populations (Figure 3F). In particular, they were not detectably expressed in luminal progenitors, which are the proposed cells of origin for basal tumors (Molyneux et al. 2010). Therefore, expression of CT83/HORMAD1 in basal tumors does not seem to merely reflect pre-existing expression in the cells of origin of the tumors.

We investigated this question further using single-cell RNA-seq data from normal human mammary glands. Using a combination of dimensional reduction, unsupervised clustering approaches, and previously known markers, we were able to separate the luminal from the basal-epithelial compartments (Figure 3G). The expression of MSRB3 and ESR1 marked the expected populations (Figure S3D). We detected some normal cells expressing CT83 and/or HORMAD1 (Figure 3G, red circles), however these cells were very rare: only 15 out of 24 292 total cells expressed HORMAD1 and/ or CT83. The positive cells either that could be assigned to a cluster were mostly “Luminal Epithelial” cluster, however more than 99% of Luminal Epithelial cells failed to express HORMAD1 or CT83, which is consistent with the lack of detection in the sorted cell populations of Figure 3F.

### Expression of HORMAD1 and CT83 in tumors correlates with promoter demethylation

Basal-like tumors are genetically unstable (Russnes et al., 2017), so we examined whether HORMAD1 and CT83 over-expression could be due to gene amplification. We found two results arguing against this possibility. First, there were no correlations between Copy Number Variation (CNV) and mRNA levels for HORMAD1 or CT83 in basal tumors (Figure 4A). Second, if the genes were overexpressed because their locus is amplified, then we would expect to see a positive correlation between the expression of HORMAD1 and its two adjoining genes (GOLPH3L, 1kb away, and CTSS, 9 kb away), and/or between CT83 and its contiguous gene SLC6A14 (250 base pairs away). We failed to detect any such correlation, whereas the expression of a gene known to undergo amplification and used as a positive control in the analysis, ERBB2, correlated positively with the expression of the neighboring gene PGAP3 (Figure 4B).

**Figure 4:**
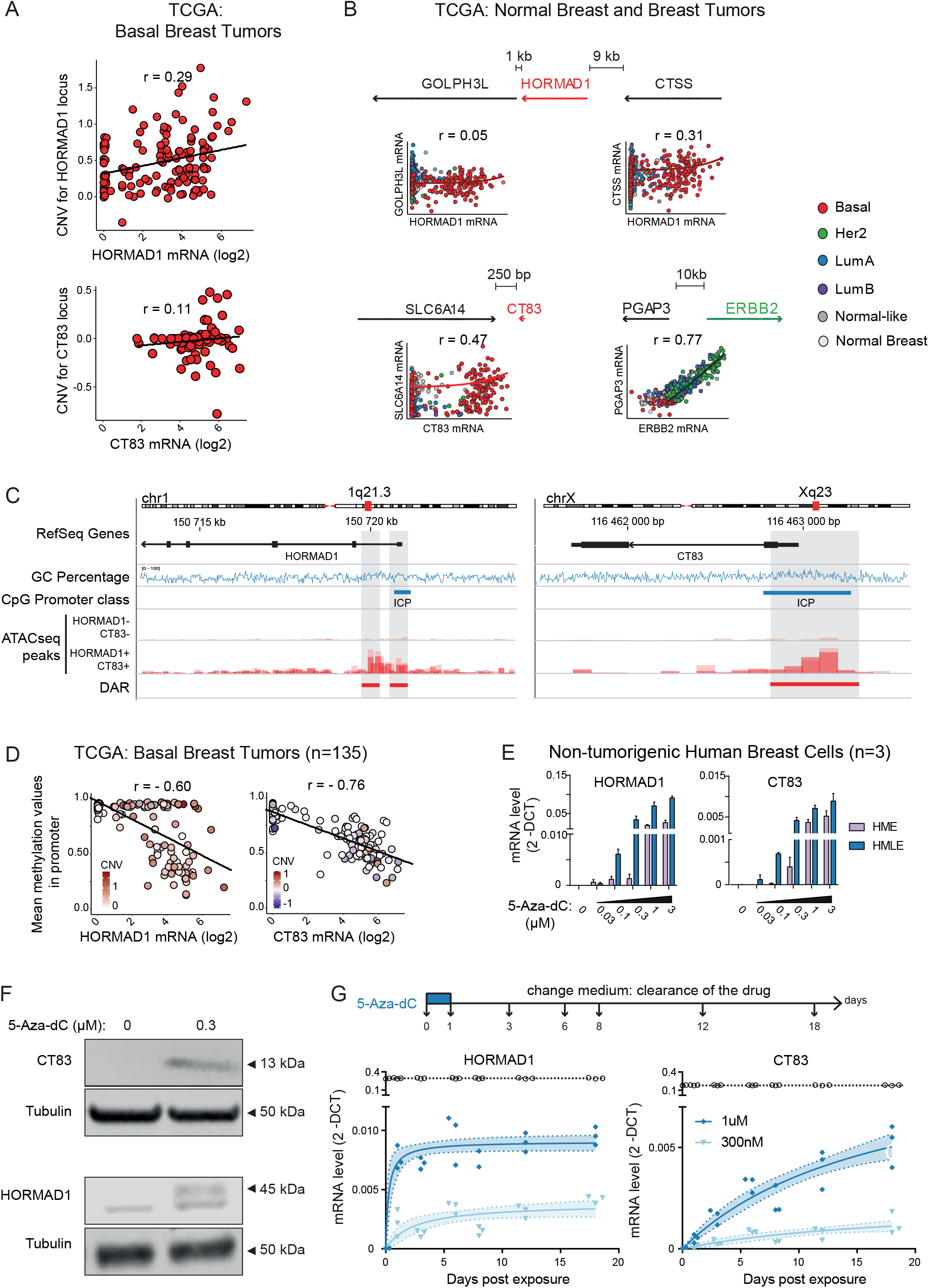
HORMAD1 & CT83 expressions are epigenetically regulated, with an essential contribution of DNA methylation. A. Correlation between HORMAD1 and CT83 expression and mean DNA methylation of their promoters (TSS +/- 200bp), according to the copy number variation of their genomic loci. B. Correlation between HORMAD1 or CT83 expression, and the expression of their neighboring genes. ERBB2 is a positive control. The color code corresponds to PAM50 tumor subtypes C. IGV representation of HORMAD1 and CT83 genomic loci, with CpG density promoter classification according to the Weber/Schübeler criteria (PMID: **17334365**). ATAC-seq data are from representative basal-like tumors (TCGA cohort). Differentially accessible regions (DAR) between these two groups of basal tumors were identified. D. Inverse correlation between HORMAD1 and CT83 expression and the mean DNA methylation of their promoters (TSS +/- 200bp). Each dot represents a tumor, and the color intensity indicates Copy Number Variation. E. RTqPCR analysis of HORMAD and CT83 expression in non-tumorigenic human mammary cell lines, in control condition or following a 48 hours 5-Aza-dC treatment at various concentrations. F. Western Blot of HORMAD1 and CT83 expression in non-tumorigenic human mammary cell lines, in control condition or following a 48 hours 5-Aza-dC treatment at 0.3 μM. G. RT-qPCR analysis of HORMAD and CT83 expression at various time points, in the same cell line, after an initial perturbation with 0.3 or 1 μM 5-Aza-dC followed by a recovery period in drug-free medium.

As amplification seemed unlikely to explain the overexpression of HORMAD1 and/or CT83, we next examined epigenetic events. The genes lack CpG islands, but both have promoters with an intermediate CpG density (ICP) (Figure 4C). These promoters overlap ATAC-seq peaks that are present in HORMAD1/CT83-expressing basal-like breast tumors, but absent in non-expressing tumors (Figures 4C and S4C). We next investigated the DNA methylation status of these promoters, using the Illumina 450K arrays available in TCGA and GEO. As shown in Figure S4B, we found high levels of methylation on the HORMAD1 and CT83 promoters in normal breast samples (that do not express the genes) and low levels of methylation in the sperm samples (where the genes are on). The data in tumors show a very strong correlation between expression and promoter demethylation for CT83 (Figure 4D). The correlation is present but less absolute for HORMAD1, as some tumors overexpress HORMAD1 without displaying demethylation. These specific tumors tend to have a higher HORMAD1 copy number (Figure 4D), and our hypothesis is that most of the copies are methylated and silent, while a few are demethylated and active.

We then tested functionally whether demethylation suffices to induce HORMAD1 and CT83 expression. For this, we used immortalized human mammary epithelial cells (HME and HMLE, Elenbaas et al. 2001) treated in vitro with 5-aza-deoxy-cytidine (5-aza-dC). The treatment induced both genes, in a dose-dependent manner (Figure 4E), and led to detectable protein expression (Figure 4F). Importantly, the genes remained expressed even after the drug was removed (Figure 4G), demonstrating a memory effect.

To better characterize the epigenetic landscape of HORMAD1 and CT83 in both normal and pathological conditions, we used public ChIP-seq datasets. In the testis, HORMAD1 showed a significant enrichment in the activating histone marks H3K27ac and H3K4me3, which were absent in breast. Conversely, in the breast, HORMAD1 and CT83 were marked by the repressive chromatin mark H3K9me3 (Figure S4A). The activation marks H3K27ac and H3K4me4 were also found for HORMAD1 and CT83 in the basal-like breast cancer cell line MDA-MB-436; but surprisingly we did not detect repressive marks in the non-tumorigenic mammary cell line MCF10A nor in the luminal A breast cancer cell line MCF7 (Figure S4B). From these data we conclude that HORMAD1 and CT83 are normally silenced by DNA methylation and, likely, H3K9me3 methylation, and that these marks are lost and replaced by active modifications such as H3K4me3 in cell lines and tumors that re-express the genes.

## DISCUSSION

### A new approach identifies cancer/testis genes expressed in different breast tumor subtypes

Cancer/Testis genes hold promise as markers, actors, and targets in cancer. Here we implemented a new bioinformatic approach to identify the Cancer/Testis genes that are overexpressed in breast cancer. This approach has the advantage of being rigorous and calculation-efficient, immediately usable for any tumor type, but also easily adaptable to seek other types of genes misexpressed in tumors. It complements previous approaches based on expression thresholds (Rousseaux et al. 2014) or vector colinearity (Wang et al. 2016), and yielded results that either approach alone would not have yielded (Figure 1B).

This approach, combined with machine learning on large breast cancer cohorts, has led us to uncover new markers that are specific of different breast cancer subtypes. Most of them were previously unknown, and some of them are associated with prognosis and response to treatment: they may become valuable markers. In addition, future investigations could examine whether they actively participate in the transformation process. Examination of early-stage tumors should reveal if the pattern of cancer/testis genes expression is determined early on, which will have interesting practical and conceptual implications.

We identify two genes —CT83 and HORMAD1— that are expressed by most basal tumors, but few other tumor of the other subtypes. By definition, these genes are normally expressed in the testis. HORMAD1 is expressed in pre- leptotene spermatocytes (Shin et al., 2010), it is required for the promotion of non-conservative recombination events in meiosis and the resulting formation of the synaptonemal complex (Kumar et al. 2015). CT83 (also known as CXorf61 or KK-LC-1) encodes a small protein (113 AA) of unknown function, normally expressed in mature sperm (Jung et al. 2019).

Both genes had been previously linked to basal tumors (Holm et al. 2016; Kaufmann et al. 2019; Wang et al. 2018; Watkins et al. 2015; Zhong et al. 2020), but our work goes further and brings a number of novel findings : 1) we rigorously prove that the genes are the 2 strongest predictors of a tumor being basal in independent cohorts, 2) we show that the genes are not expressed in healthy breast progenitors, showing that the induction occurs de novo.

Three important questions remain open and will be discussed briefly in the following paragraphs: what is the order of events leading to HORMAD1/CT83 induction in basal tumors? And what are the mechanistic bases for their induction?

### Order of events

About 90% of basal tumors in the TCGA cohort express HORMAD1 or CT83, and about 60% express both. There are two non-exclusive interpretations for these high proportions.

First, the induction of the genes could be an early event that occurs in most early lesions and is maintained as the tumor progresses. In principle, this deregulation could even occur earlier than the main transforming event, such as activation of Myc. It could be that HORMAD1/CT83 induction reflects a disturbed epigenetic landscape in rare tumor-initiating cells, which could itself increase the probability of cellular transformation. In that possibility, HORMAD1 and CT83 themselves could just be markers of the early epigenetic instability, or they could actively participate in the ensuing transformation. One piece of data supporting this “induction before transformation” hypothesis is that a few rare cells in the healthy breast already express CT83 and/or HORMAD1. Some of those aberrant cells might eventually be amenable to enter the basal-like transformation path.

Second, it could be that the expression of both HORMAD1 and CT83 occurs after transformation and brings a selective advantage to basal tumor cells. The genes have only been studied individually so far, but there is convincing evidence that HORMAD1 overexpression impairs homologous recombination and increases genomic instability in basal breast tumor cells, therefore possibly speeding up tumor evolution (Watkins et al. 2015). HORMAD1 overexpression is also detected in lung tumors but, paradoxically, it seems to increase the robustness of homologous recombination in these tumors, making them more resistant to DNA-damaging chemotherapy. These divergences may mean that HORMAD1 has context-dependent functions, for instance in the presence or absence of other actors such as CT83.

### Mechanism of induction

While basal tumors are genetically unstable, we rule out gene amplification as the main mechanism of HORMAD1/ CT83 induction. Instead, we show that DNA methylation is a barrier to HORMAD1/CT83 activation, which is consistent with previously published reports (Nichols et al. 2018; Wang et al. 2018; Chen et al. 2019). Importantly, we find that, once the genes have been induced by a 5-aza-deoxycytidine treatment, they remain active even when 5-aza-dC has been removed. In other words, they switch to a stable “On” state. This makes them excellent markers of past epigenetic disturbances.

Further investigations will be required to elucidate the initial event(s) that lead to the derepression of HORMAD1/CT83 at some point during the history of most basal tumors. It could be a stochastic phenomenon occuring before or after transformation; alternatively it could be a directed event triggered by the transforming pathway(s). At any rate, many cancer/testis genes are repressed by DNA methylation, but HORMAD1 and CT83 are highly specific in their association with basal tumors, so they could be specifically induced in this tumor type, specifically selected for, or both.

### Limits and perspectives

We note that our analysis has a number of possible limitations. One is that we used pre-existing lists of cancer/testis genes; any gene not detected in these previous publications has not been considered in our work. Another has to do with sensitivity: if certain genes are expressed only in a small number of tumors, then the smoothing we performed in the initial step of our analysis may have made them undetectable. Our sample size was large, with more than 1000 tumors, but certain rare subtypes (such as normal-like tumors, only represented by 40 data points) may benefit from a more focused approach. Also, we focused on one specific type of genes misexpressed in tumors: the cancer/testis genes. However, other tissue-specific genes ectopically expressed in breast tumors can be a rich source of markers and may be involved in the transformation process. These genes can be easily recovered from our dataset and may deserve further investigations in the future.

In spite of the limitations mentioned above, the current work brings new conceptual insight into the role of cancer/ testis genes in breast cancer, showing that reactivation occurs de novo and could have a synergistic effect. In practical terms, as already underlined by other investigators, the genes we have studied represent potential targets for immunotherapy. We show, in addition, that their epigenetic activation seems irreversible, and that they could constitute ideal witnesses of past episodes of epigenetic instability. This may help better understand the role of epigenetic instability in breast tumors, and its mechanistic connection to cellular transformation.

## MATERIEL & METHODS

### Wet biology

#### Cell culture

Human mammary cell lines, derived from normal mammary tissue, were obtained from collections developed and generously given by the laboratories of Christophe Ginestier (CRCM) and Raphaël Margueron (Institut Curie). Cancer cell lines (MDA-MB-436, HEK293T) were obtained from ATCC or generously given by the laboratory of Marc-Henri Stern (Institut Curie).

The cell lines were grown using the recommended culture conditions. Cells were incubated in a humidified atmosphere at 37°C under 5% CO2. All experiments were done with subconfluent cells in the exponential phase of growth.

| Cell lines | Medium |
| --- | --- |
| HME, HMLE | DMEM:F12 medium supplemented with 10% FBS, 1% penicillin/streptomycin, Non-Essential Amino Acids (LifeTechnology 11140-035) 1%, Insulin Humalog (Lily) 10ug/ml, Hydrocortison (Serb) 0.5 ug/ml, EGF (ThermoFisher PHG0311) 10ng/ml |
| HEK293T, MDA-MB-436 | DMEM medium supplemented with 10% FBS, 1% penicillin/streptomycin |

#### Treatment of cells with 5-aza-dC

Treatment with 5-Aza-dC was performed as described previously (Naciri et al. 2019). Briefly, for dose-response experiments, cells were seeded at a density of 1. 10^4^ cells in a 6-well tissue culture plate. When cells became firmly adherent to plastic, the medium was replaced with fresh medium containing the appropriate concentration of 5-Aza-dC, every 24h for 2 days (two pulses). For the recovery assay, cells were seeded at a density of XXX in a 100 mm tissue culture plate. When cells became firmly adherent to plastic (T0), the medium was replaced with fresh medium containing 1 uM or 300 nM of 5-Aza-dC for 24h (one pulse). At the end of the treatment, the medium was replaced with fresh culture medium without 5-Aza-dC, and cells were cultured for an additional 2 weeks in subconfluent condition with regular passages. At the end of the treatment and at the appropriate time-points, cells were used for molecular assays. Control cultures were treated under similar experimental conditions in the absence of 5-Aza-dC.

#### Generation of the HORMAD1 and/or CT83 mammary cell lines

The maximal reporter cassette comprised HORMAD1-P2A-CT83-T2A-Blasti^R^ (Synthesized by GenScript). The three proteins expressed by the cassette were separated from each other by self-cleaving 2A peptides (P2A, T2A). This cassette was cloned in a lentiviral backbone from ORIGENE (derived from PS100071), under the control of the constitutive CMV promoter. The control plasmid (Blasti^R^) and the two other plasmids (HORMAD1 -T2A-Blasti^R^ and CT83-T2A-Blasti^R^) were generated by enzymatic digestion; all the plasmids were grown and prepared individually. The sequences were validated by sequencing. Lentiviruses were generated and used for transduction. Production of lentiviral particles was performed by calcium-phosphate transfection of HEK293T with psPAX2 and pMD2.G plasmids, in a BSL3 tissue culture facility. HME or HMLE cells were seeded into 12-well plates, infected, and selected with blasticidin (5ug/ml) for 15 days.

#### Western blotting

Cells were harvested and lysed in RIPA buffer (Sigma) with with protease inhibitor cocktail (Thermo Fisher Scientific), sonicated with a series of 30s ON / 30s OFF for 5 min on a Bioruptor (Diagenode), and centrifuged at 16,000 g for 5 min at 4°C. The supernatant was collected and quantified by BCA assay (Thermo Fisher Scientific). Thirty microgram protein extract per sample was mixed with NuPage 4X LDS Sample Buffer and 10X Sample Reducing Agent (Thermo Fisher Scientific) and denatured at 95°C for 5 min. Samples were resolved on a pre-cast SDS-PAGE 4-12% gradient gel (Thermo Fisher Scientific) with 120V electrophoresis for 90 min and blotted onto a nitrocellulose membrane (Millipore). The membrane was blocked with 5% fat-free milk/PBS at RT for 1 h, then incubated overnight at 4°C with appropriate primary antibodies. After three washes with PBS/0.1% Tween20, the membranes were incubated with the cognate fluorescent secondary antibodies and revealed in the LI-COR Odyssey imaging system. The following antibodies were used in this study: α-HORMAD1 (dilution 1:1000, reference HPA037850), α-CT83 (dilution 1:1000, reference HPA004773), α-Tubulin (dilution 1:10 000, reference Abcam ab7291).

#### Quantitative Real-time PCR

RNA extraction was doing using Tri reagent according to the manufacturer’s recommendations. One microgram of total RNA was reverse transcribed using SuperScript IV Reverse Transcriptase (Thermo Fisher Scientific) and Oligo dT primers (Promega). qPCR was performed using Power SYBR Green (Applied Biosystems) on a Viia 7 Real-Time PCR System (Life Tech). *TBP* and *PGK1* genes were used for normalization of expression values. Primer sequences are available in Supplementary Table S4.

### Bioinformatics

#### Public data sets used in this study

We used previously published gene lists to define testis-specific genes, tumor suppressor genes and oncogenes. We also used multiple public datasets involving both normal and tumor tissues to evaluate C/T gene expression. Detailed information of these databases was listed in the Supplementary Table 5.

#### Development of the Cancer-Gene Markers Detection pipeline

Briefly, we computed the Kernel’s density estimation for each gene expression pattern in healthy mammary gland and in breast cancer cohorts, respectively. We then analyzed density profiles variations using the derivative of the density functions, and classify genes as unimodal or multimodal in normal mammary tissues and breast cancer samples. For each gene, we calculated the mean expression values in normal and cancers samples. We classify genes according to these parameters, as described in figure S1A-C. All the detailed scripts are available on GitHub (https://github.com/MartheLaisne/CTA_BreastCancers).

#### Identification of genes with abnormal breast cancer expression pattern using transcriptomic TCGA analysis

TCGA gene count datasets for breast normal and cancer samples were downloaded using TCGAbiolinks (Colaprico et al., v2.10.5). Expression were normalized with DESeq2 (Love MI, Huber W, Anders S, v.1.22.2). Abnormally expressed genes were defined as any expression value greater than the mean expression + 3 standard-deviations in normal mammary tissues. All the detailed scripts are available on GitHub.

#### Validation of the Testis-specific expression pattern for the selected 139 C/T genes

Expression values for GTEx (Carithers LJ et al., 2015) dataset was obtained directly from the project webpage as TPM values, and the median expression values by tissue were calculated. We extracted expression values for the 139 selected TS genes, and we performed an unsupervised clustering (Euclidean distance and complete method) of the genes and the samples based on these values. Detailed script is on GitHub.

#### Analyze of the INVADE dataset

Briefly, raw counts were normalized using DESeq2 (Love MI, Huber W, Anders S, v.1.22.2). because there are no normal tissues in this dataset, another strategy was used to defined the threshold for abnormal C/T gene activation: we used the bimodality of the expression values distribution to define a background level. Any expression value below this threshold was considered as noise, and the gene as repressed. The top 20 CT genes based on random forest analyzes were used to performed an unsupervised hierarchical clustering (binary distrance and Ward.D2 method) of the 55 tumors samples. Detailed script is on GitHub.

#### Survival and drug-response analysis

For recidive-free survival (RFS), data were download from https://kmplot.com/analysis/ (n=4934), using the indicated parameters for sample selection. Data were then analyzed using custom R script and surviminer and survival R packages. For anthracyclin-response analysis, data were dowload from http://www.rocplot.org/ using the indicated parameters for sample selection, and analyzed using standard R functions. ROC curves were generated using ROCit R package.

#### Analyze of normal mammary breast microarray

Data were download at https://www.ebi.ac.uk/arrayexpress/experiments/E-MTAB-4145. The raw CEL data were normalized using the following packages: affy (v1.60.0), ArrayExpress (v1.42.0) for annotation and data importation; oligo (v1.45.0), arrayQualityMetric (v3.38.0) for quality control and pre-processing; limma (v3.38.3) for analysis and statistics.

#### scRNAseq of normal mammary breast cells

Briefly, data were download (GSE113197) and analyze using Seurat (v3.1.4) package. For the normalization, we keep unexpressed genes because we are interested in C/T genes, which are expected to not be expressed in healthy mammary cells. We filtered cells to keep only cell with at least 500 genes detected, but no more than 6000, and less than 10% of mitochondrial gene expressed. UMAP was performed using the 10 first components of the PCA. Cell identities were assigned based on the expression of lineage markers (source code is at: https://github.com/Michorlab/tnbc_scrnaseq/blob/master/code/funcs_markers.R). Detailed script is on GitHub.

#### scRNAseq of triple-negative breast tumors

FASTQ read pairs were aligned to the human reference genome (build gencode v29) using STAR (v2.7.5c) and default single-pass parameters. Uniquely aligned reads were kept for downstream analysis using Samtools view (v1.10) and parameters: -q 10 -b –o, and counted with htseq (--stranded=yes –type=exon). Data were analyzed using Seurat (v3.1.4). As for Healthy mammary scRNAseq analyze, we identified low quality cells by (i) few expressed genes, (iii) abnormally high number of expressed genes and (iii) high mitochondrial gene expression. Cell identities were determined using the same procedure than for the healhy mammary scRNAseq data. We also used Lehman signature to assigned each cancer cell to a lehman subtype, as described in the original publication (code source: https://github.com/Michorlab/tnbc_scrnaseq)

#### Differential Gene Expression Analysis in TCGA basal-like samples

HORMAD1- and CT83-positive tumors were identified based on normalized RNAseq (FPKM-UQ) data downloaded from TCGA (2020 accession). Briefly, we defined a threshold for positive HORMAD1 and CT83 expression based on the expression level detected in non-tumor breast samples (NT) as follow:

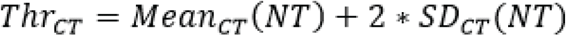

We classified tumors in 4 different groups based on their expression levels of both HORMAD1 and CT83. Then, we download HTseq-counts data for basal-like breast tumors only and we performed a differential expression analysis using the R package *DESeq2*, with the HORMAD1 & CT83 label as factor of interest. Differentially expressed genes were defined with p-adjusted < 0.05 and absolute value for the fold-change > 1.5.

#### Differential Peaks Intensity Analysis in TCGA basal-like samples

Both raw counts ATAC-seq data and gene expression data from TCGA were accessed (2020 accession) through either the Genomic Data Commons (GDC) using the GDC Data Transfer Tool Client or the data transfer tool TCGAbiolinks (Colaprico 2016). Individual patient files were assembled using in-house scripts in an R computing environment. Preprocessing consisted of patient and gene matching between data types, log transformation of gene expression data, and classification of the ATAC-seq samples regarding to their HORMAD1 / CT83 expression status, defined in the previous section. For differential analysis, we basal-like tumors from ATAC-seq datas (n=30). Differential peak intensities were found using *DESeq2*. Differentially open regions were defined with p-adjusted < 0.01 and absolute value for the fold-change > 2.

#### CpG promoter classes identification

Promoters were according to the hg38 version of the human genome, as described in the original article (Weber et al. 2007). Briefly, promoters were classified in three categories to distinguish strong CpG islands, weak CpG islands and sequences with no local enrichment of CpGs. We determined the GC content and the ratio of observed versus expected CpG dinucleotides in sliding 500-bp windows with 5-bp offset. The CpG ratio was calculated using the following formula: (number of CpGs × number of bp) / (number of Cs × number of Gs). The three categories of promoters were determined as follows: HCPs (high-CpG promoters) contain a 500-bp area with CpG ratio above 0.75 and GC content above 55%; LCPs (low-CpG promoters) do not contain a 500-bp area with a CpG ratio above 0.48; and ICPs (intermediate CpG promoters) are neither HCPs nor LCPs.

#### Correlation DNA methylation data and expression data for TCGA samples

Both DNA methylation data, Copy Number Variations (CNV) data and gene expression data from TCGA were accessed (2020 accession) through either the Genomic Data Commons (GDC) using the GDC Data Transfer Tool Client or the data transfer tool TCGAbiolinks (Colaprico 2016). Individual patient files were assembled using in-house scripts in an R computing environment. Preprocessing consisted of patient and gene matching between data types and log transformation of gene expression data. The methylation data in this study were acquired by the Illumina 450K array, which interrogates more than 450 000 methylation sites on the Illumina chip. The data for this study contained information of 485 578 CpG sites. The CNV data were acquired by the Affymetrix SNP 6.0 array numeric CNV values were derived from GISTIC2.

Correlation analysis was performed using Pearson’s correlation. The correlation was performed between methylation beta values (respectively between CNV values) and log-base-2-transformed gene expression data with a *p*-value threshold of 0.05. All statistical tests used standard R functions.

#### Correlation adjacent genes TCGA

Correlation analysis was performed using Pearson’s correlation. The correlation was performed between the two log2 normalized adjacent genes expression values. All statistical tests used standard R functions.

## ACKNOWLEDGEMENTS

We are grateful to the following colleagues for helpful discussions: Saadi Khochbin, Fatima Mechta-Grigoriou, Marc-Henri Stern, Raphael Margueron, Laia Richart, Céline Vallot, Josh Waterfall, Julie Cocquet, Valérie Borde, Cathy Jackson, Jafar Sharif, Julien Sage, Bernard de Massy.

## Plateformes

Cytométrie IJM, Epigénétique EDC, Microscopie IJM/EDC

## Figure Legends

**Figure S1:**
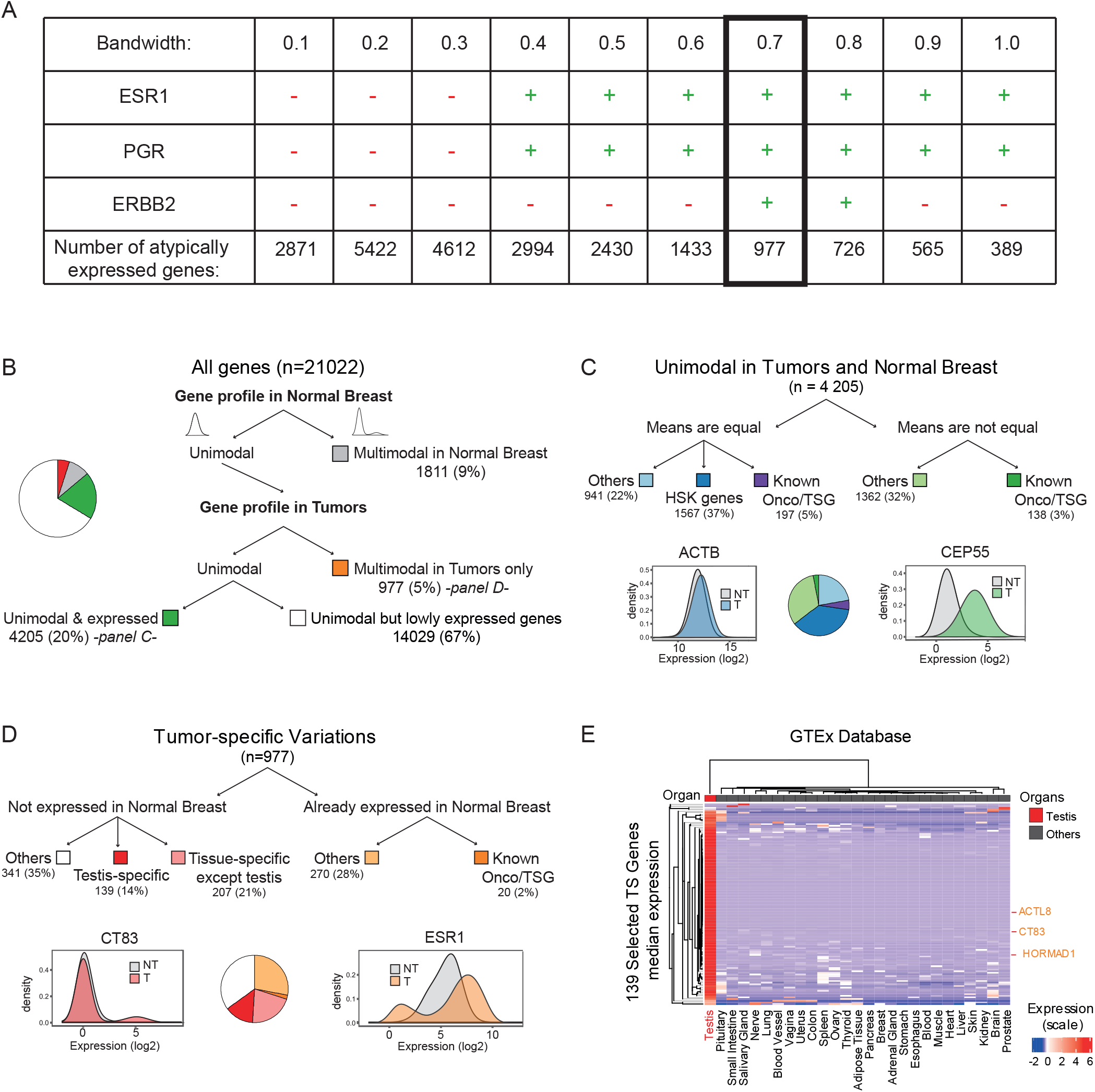
Optimizing parameters for the bioinformatic screen, further uses and validations. A. Outputs of the screen for different smoothing parameters (Bandwidth). Previously known breast cancer markers (*ESR1, PGR, ERBB2*) were used as positive controls, and housekeeping genes (*ACTB, GAPDH, TUBA1A*) were used as negative control. A red minus sign means the gene was not detected as aberrantly expressed in tumors, a green plus sign means that it was. The total number of atypically expressed genes for each bandwidth value is shown. B. Classification of all genes according to our parameters: we were interested in genes with a homogeneous expression in NB (*ie*. Unimodal profile in NB). Then, these genes can be subsequently divided according to their expression pattern in breast tumors: two situations were of specific interest: genes that are homogeneously expressed in breast tumors too (panel C), and genes that are overexpressed or repressed in a subset of breast tumors (panel D). C. Refinement of the characterization of homogenously expressed genes in NB and in breast tumors, respectively: when means were significantly different in NB and in Tum, these genes could be used as tumor markers. Some of such genes are known overexpressed oncogenes or repressed tumor suppressor genes; a significant part of them (1362 genes) are unknown but could play a role in breast tumor development D. Refinement of the characterization for tumor-specific variables genes: approximatively 70% of them are repressed in NB and abnormally activated in breast tumors; amongst these genes there are known tissue-specific genes (including testis-specific genes). The remaining 30% are overexpressed or repressed genes in some breast tumors, including known subtype-specific oncogenes like *ESR1*, and others genes that could be used as marker of specific tumor subgroups. E. Heatmap showing the mean expression values (Z-score) for the 139 selected C/T genes in various human adult tissues, based on RNA-seq data from GTEx.

**Figure S2.**
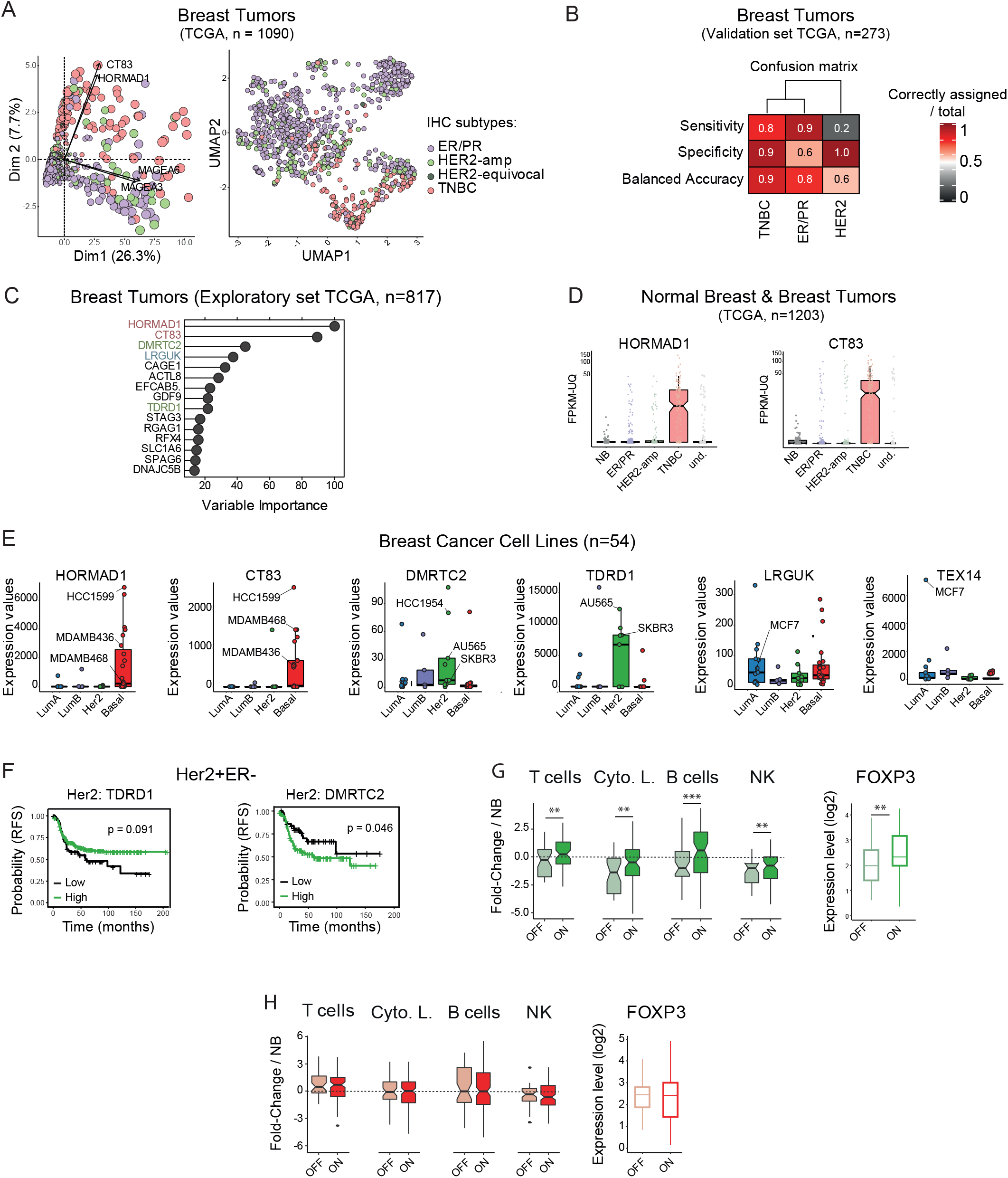
Examination of marker expression in tumors classified by IHC and in tumor cell lines. Expression in early tumors and association with survival. A. Multidimensional analysis of TCGA breast tumor and healthy samples based on the 139 selected C/T gene expression. Each dot is a sample, color code corresponds to immunohistochemistry (IHC) classification (based on ER/PR/HER2 expression). Left: Principal Component Analysis, dot sizes are proportional to the quality of representation in PC1/PC2 space. The best correlated C/T genes to PC1/PC2 are represented. Right: Uniform Manifold Approximation and Projection B. Confusion matrix for breast tumor samples in the validation cohort (randomly selected 25% samples from the TCGA breast tumors) of the IHCtumor subtypes prediction obtained with the best Random Forest model. This model was established after a 500 trees training on the discovery cohort (the remaining 75%), based on the expression level of the 139 C/T C. Top 15 most important variables in the best Random Forest model for IHC tumor subtype prediction. D. Expression levels for the 2 basal-specific C/T genes in the breast TCGA cohort, according to IHC tumor subtype E. Expression levels for 6 subtype-specific C/T genes in breast cancer cell lines from the Cancer Cell Line Encyclopedia, according to PAM50 tumor subtype. Some commonly used cell lines are highlighted. F. Relapse-free survival curves for Her2-positive breast cancer patients, according to DMRTC2 expression (left), or TDRD1 expression (right). G. Immune infiltration of Her2-positive breast tumors that express (ON) or do not express (OFF) DMRTC2, inferred from whole tumor RNA-seq data using MCPcounter. Fold-Change were computed against Normal Breast (NB). Right: Expression level of the immune suppressive factor *FOXP3* in the same tumors. P-value < 0.01: ** ; P- value < 0.001: *** H. Same as panel H, but for basal-like tumors that either express (ON) or do not express (OFF) HORMAD1 and CT83.

**Figure S3:**
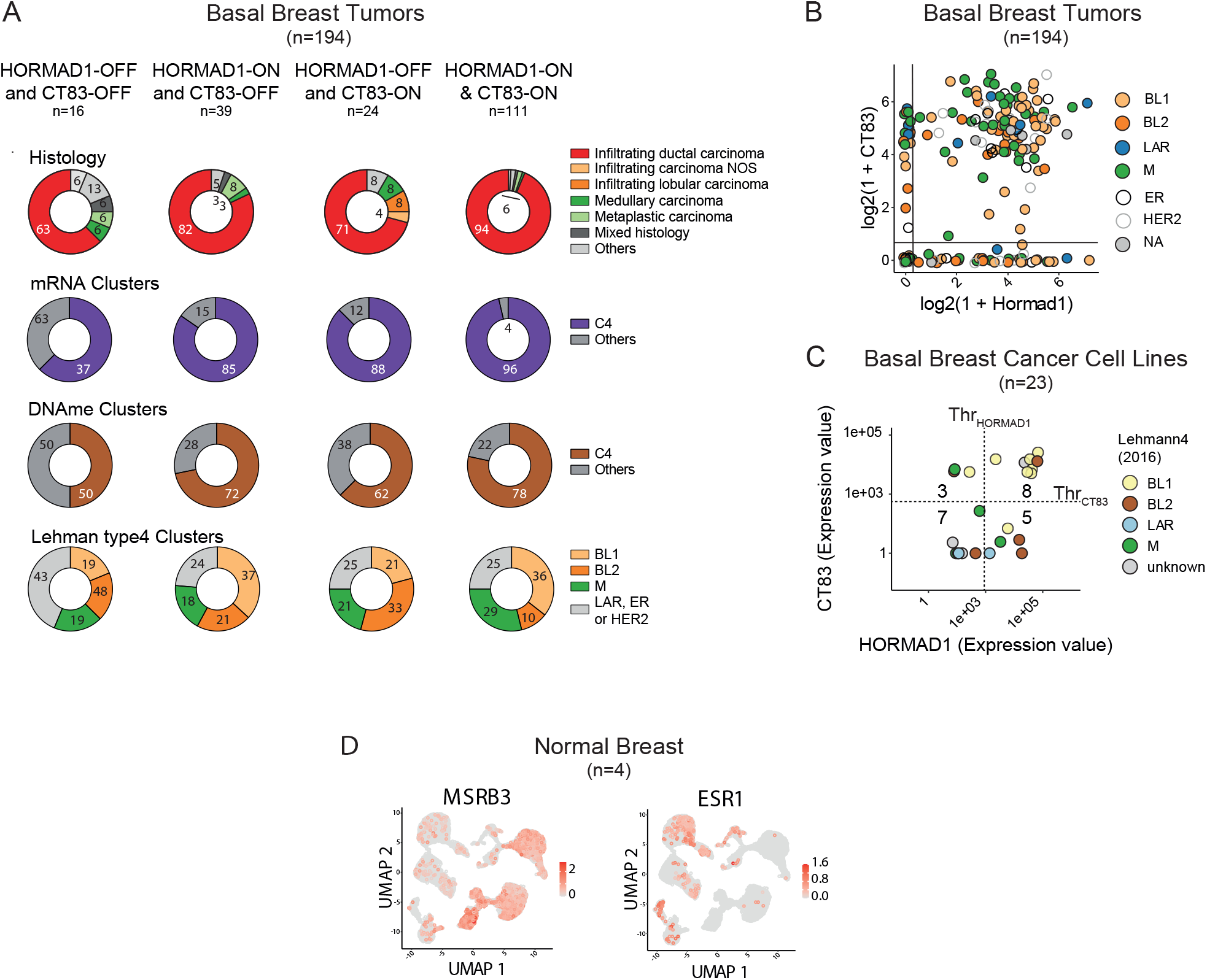
Characteristics of HORMAD1/CT83-positive basal tumors and cell lines, validation by IHC. A. Characteristics of basal-like breast tumors from the TCGA according to their activation status of HORMAD1 and CT83. B. Links between HORMAD1 and CT83 expression and Lehman’s basal tumor subgroups C. Co-expression of *HORMAD1* and *CT83* based on RNA-seq analysis (log2 Normalized expression) in basal-like breast cancer cell lines (n= 22) from the CCLE. Thresholds were calculated as in Fig. 3A. D. UMAP representation of a scRNA-seq study on 4 healthy mammary glands (GSE113197), after an enrichment in epithelial cell by FACS. MSRB3 or ESR1 expression marks the expected populations.

**Figure S4:**
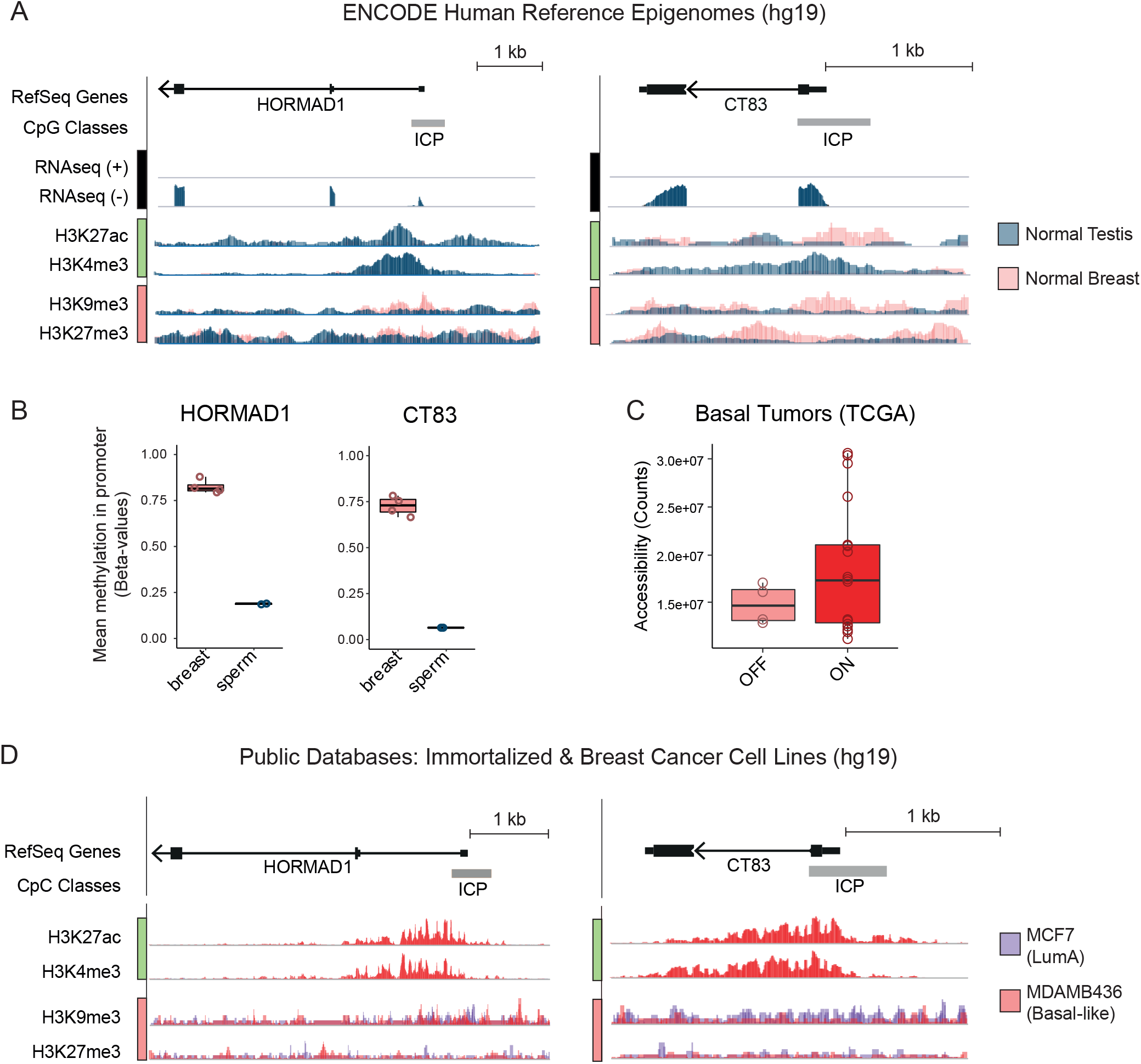
Epigenetic landscapes of the HORMAD1 and CT83 genes. A. IGV representation of transcriptomic and histone modification landscapes at HORMAD1 and CT83 loci in healthy Testis and Breast samples (ENCODE). B. DNA methylation levels on the promoter of HORMAD1 or CT83, in normal human breast and sperm, from 450K array values. C. Total accessibility scores for HORMAD1- and CT83-negative or positive basal-like TCGA tumors, calculated based on ATAC-seq data. D. IGV representation of histone modifications landscapes at HORMAD1 and CT83 loci in breast cell lines: MCF7 cells do not express HORMAD1 or CT83, whereas MDA-MB436 cells express both.

**TS1:** 139 Selected Cancer/Testis genes

| gene_id | gene_id | entrezgene | ensembl_ |
| --- | --- | --- | --- |
| CRISP1\|167 | CRISP1 | 167 | ENSG00000124812 |
| AQP5\|362 | AQP5 | 362 | ENSG00000161798 |
| CSH2\|1443 | CSH2 | 1443 | ENSG00000213218 |
| CTAG1B\|1485 | CTAG1A1 | 485 | ENSG00000183678 |
| CTAG1B\|1485.1 | CTAG1B1 | 485 | ENSG00000184033 |
| CTNNA2\|1496 | CTNNA2 | 1496 | ENSG00000066032 |
| CYP19A1\|1588 | CYP19A1 | 1588 | ENSG00000137869 |
| DMP1\|1758 | DMP1 | 1758 | ENSG00000152592 |
| GDF9\|2661 | GDF9 | 2661 | ENSG00000164404 |
| GIP\|2695 | GIP | 2695 | ENSG00000159224 |
| GPX5\|2880 | GPX5 | 2880 | ENSG00000224586 |
| HSD3B1\|3283 | HSD3B1 | 3283 | ENSG00000203857 |
| IGFBP1\|3484 | IGFBP1 | 3484 | ENSG00000146678 |
| INSL3\|3640 | INSL3 | 3640 | ENSG00000248099 |
| INSL4\|3641 | INSL4 | 3641 | ENSG00000120211 |
| KRT33B\|3884 | KRT33B | 3884 | ENSG00000131738 |
| MAGEA1\|4100 | MAGEA1 | 4100 | ENSG00000198681 |
| MAGEA2\|4101 | MAGEA2 | 4101 | ENSG00000184750 |
| MAGEA2\|4101.1 | MAGEA2B | 4101 | ENSG00000183305 |
| MAGEA3\|4102 | MAGEA3 | 4102 | ENSG00000221867 |
| MAGEA6\|4105 | MAGEA3 | 4105 | ENSG00000221867 |
| MAGEA6\|4105.1 | MAGEA6 | 4105 | ENSG00000197172 |
| MAGEA8\|4107 | MAGEA8 | 4107 | ENSG00000156009 |
| MAGEA10\|4109 | MAGEA10 | 4109 | ENSG00000124260 |
| MAGEA12\|4111 | MAGEA12 | 4111 | ENSG00000213401 |
| MAGEB2\|4113 | MAGEB2 | 4113 | ENSG00000099399 |
| NMBR\|4829 | NMBR | 4829 | ENSG00000135577 |
| PENK\|5179 | PENK | 5179 | ENSG00000181195 |
| PSG9\|5678 | PSG9 | 5678 | ENSG00000183668 |
| PSG11\|5680 | PSG11 | 5680 | ENSG00000243130 |
| PSG11\|5680.1 | PSG3 | 5680 | ENSG00000221826 |
| RFX4\|5992 | RFX4 | 5992 | ENSG00000111783 |
| SLC1A6\|6511 | SLC1A6 | 6511 | ENSG00000105143 |
| SSX1\|6756 | SSX1 | 6756 | ENSG00000126752 |
| SSX5\|6758 | SSX5 | 6758 | ENSG00000165583 |
| AURKC\|6795 | AURKC | 6795 | ENSG00000105146 |
| TNP1\|7141 | TNP1 | 7141 | ENSG00000118245 |
| DNALI1\|7802 | DNALI1 | 7802 | ENSG00000163879 |
| TKTL1\|8277 | TKTL1 | 8277 | ENSG00000007350 |
| PAGE1\|8712 | PAGE1 | 8712 | ENSG00000068985 |
| XAGE2\|9502 | XAGE2B | 9502 | ENSG00000155622 |
| XAGE2\|9502.1 | XAGE2 | 9502 | ENSG00000185751 |
| XAGE1D\|9503 | XAGE1A | 9503 | ENSG00000204379 |
| XAGE1D\|9503.1 | XAGE1E | 9503 | ENSG00000204375 |
| XAGE1D\|9503.2 | XAGE1D | 9503 | ENSG00000204376 |
| XAGE1D\|9503.3 | XAGE1C | 9503 | ENSG00000183461 |
| XAGE1D\|9503.4 | XAGE1B | 9503 | ENSG00000204382 |
| PAGE4\|9506 | PAGE4 | 9506 | ENSG00000101951 |
| SPAG6\|9576 | SPAG6 | 9576 | ENSG00000077327 |
| SSX3\|10214 | SSX3 | 10214 | ENSG00000165584 |
| SSX3\|10214.1 | SSX5 | 10214 | ENSG00000165583 |
| STAG3\|10734 | STAG3 | 10734 | ENSG00000066923 |
| CAPN11\|11131 | CAPN11 | 11131 | ENSG00000137225 |
| SPO11\|23626 | SPO11 | 23626 | ENSG00000054796 |
| TMEFF2\|23671 | TMEFF2 | 23671 | ENSG00000144339 |
| AIPL1\|23746 | AIPL1 | 23746 | ENSG00000129221 |
| CABYR\|26256 | CABYR | 26256 | ENSG00000154040 |
| ZBTB32\|27033 | ZBTB32 | 27033 | ENSG00000011590 |
| RBMXL2\|27288 | RBMXL2 | 27288 | ENSG00000170748 |
| VCX2\|51480 | VCX2 | 51480 | ENSG00000177504 |
| VCX3A\|51481 | VCX3A | 51481 | ENSG00000169059 |
| L1TD1\|54596 | L1TD1 | 54596 | ENSG00000240563 |
| NXF2\|56001 | NXF2 | 56001 | ENSG00000185554 |
| NXF2\|56001.1 | NXF2B | 56001 | ENSG00000185945 |
| TEX14\|56155 | TEX14 | 56155 | ENSG00000121101 |
| TEX11\|56159 | TEX11 | 56159 | ENSG00000120498 |
| TDRD1\|56165 | TDRD1 | 56165 | ENSG00000095627 |
| ANKRD7\|56311 | ANKRD7 | 56311 | ENSG00000106013 |
| TRIM49\|57093 | TRIM49 | 57093 | ENSG00000168930 |
| SPINLW1\|57119 | EPPIN | 57119 | ENSG00000101448 |
| RGAG1\|57529 | RGAG1 | 57529 | ENSG00000243978 |
| DMRTC2\|63946 | DMRTC2 | 63946 | ENSG00000142025 |
| NEUROG2\|63973 | NEUROG2 | 63973 | ENSG00000178403 |
| EDDM3B\|64184 | EDDM3B | 64184 | ENSG00000181552 |
| C19orf57\|79173 | C19orf57 | 79173 | ENSG00000132016 |
| BCL2L14\|79370 | BCL2L14 | 79370 | ENSG00000121380 |
| LIN28A\|79727 | LIN28A | 79727 | ENSG00000131914 |
| LIN28A\|79727.1 | LIN28AP1 | 79727 | ENSG00000213120 |
| ACTL8\|81569 | ACTL8 | 81569 | ENSG00000117148 |
| TEX101\|83639 | TEX101 | 83639 | ENSG00000131126 |
| HORMAD1\|84072 | HORMAD1 | 84072 | ENSG00000143452 |
| DSCR8\|84677 | DSCR8 | 84677 | ENSG00000198054 |
| NAA11\|84779 | NAA11 | 84779 | ENSG00000156269 |
| MAEL\|84944 | MAEL | 84944 | ENSG00000143194 |
| DNAJC5B\|85479 | DNAJC5B | 85479 | ENSG00000147570 |
| FATE1\|89885 | FATE1 | 89885 | ENSG00000147378 |
| PAGE5\|90737 | PAGE5 | 90737 | ENSG00000158639 |
| TDRD12\|91646 | TDRD12 | 91646 | ENSG00000173809 |
| SYCE1\|93426 | SYCE1 | 93426 | ENSG00000171772 |
| CGB5\|93659 | CGB5 | 93659 | ENSG00000189052 |
| CGB5\|93659.1 | CGB | 93659 | ENSG00000104827 |
| CGB5\|93659.2 | CGB8 | 93659 | ENSG00000213030 |
| PNMA5\|114824 | PNMA5 | 114824 | ENSG00000198883 |
| CATSPER1\|117144 | CATSPER1 | 117144 | ENSG00000175294 |
| ZPBP2\|124626 | ZPBP2 | 124626 | ENSG00000186075 |
| C17orf64\|124773 | C17orf64 | 124773 | ENSG00000141371 |
| ZDHHC19\|131540 | ZDHHC19 | 131540 | ENSG00000163958 |
| ZFP42\|132625 | ZFP42 | 132625 | ENSG00000179059 |
| NOBOX\|135935 | NOBOX | 135935 | ENSG00000106410 |
| LRGUK\|136332 | LRGUK | 136332 | ENSG00000155530 |
| DCAF12L1\|139170 | DCAF12L1 | 139170 | ENSG00000198889 |
| MAGEB16\|139604 | MAGEB16 | 139604 | ENSG00000189023 |
| RPL10L\|140801 | RPL10L | 140801 | ENSG00000165496 |
| C20orf152\|140894 | CNBD2 | 140894 | ENSG00000149646 |
| C10orf82\|143379 | C10orf82 | 143379 | ENSG00000165863 |
| LYPD4\|147719 | LYPD4 | 147719 | ENSG00000183103 |
| FAM187B\|148109 | FAM187B | 148109 | ENSG00000177558 |
| PNLDC1\|154197 | PNLDC1 | 154197 | ENSG00000146453 |
| CSAG1\|158511 | CSAG1 | 158511 | ENSG00000198930 |
| FMR1NB\|158521 | FMR1NB | 158521 | ENSG00000176988 |
| FSIP1\|161835 | FSIP1 | 161835 | ENSG00000150667 |
| ADAD2\|161931 | ADAD2 | 161931 | ENSG00000140955 |
| RNF133\|168433 | RNF133 | 168433 | ENSG00000188050 |
| XAGE3\|170626 | XAGE3 | 170626 | ENSG00000171402 |
| XAGE5\|170627 X | AGE5 | 170627 | ENSG00000171405 |
| COX7B2\|170712 | COX7B2 | 170712 | ENSG00000170516 |
| FAM9C\|171484 | FAM9C | 171484 | ENSG00000187268 |
| SPAG17\|200162 | SPAG17 | 200162 | ENSG00000155761 |
| CXorf61\|203413 | CT83 | 203413 | ENSG00000204019 |
| PAGE2\|203569 | PAGE2 | 203569 | ENSG00000234068 |
| C18orf20\|221241 | LINC00305 | 221241 | ENSG00000179676 |
| C16orf73\|254528 | MEIOB | 254528 | ENSG00000162039 |
| WFDC11\|259239 | WFDC11 | 259239 | ENSG00000180083 |
| ODF3L2\|284451 | ODF3L2 | 284451 | ENSG00000181781 |
| CAGE1\|285782 | CAGE1 | 285782 | ENSG00000164304 |
| TMEM95\|339168 | TMEM95 | 339168 | ENSG00000182896 |
| COX8C\|341947 | COX8C | 341947 | ENSG00000187581 |
| GNAT3\|346562 | GNAT3 | 346562 | ENSG00000214415 |
| CXorf66\|347487 | CXorf66 | 347487 | ENSG00000203933 |
| C12orf42\|374470 | C12orf42 | 374470 | ENSG00000179088 |
| EFCAB5\|374786 | EFCAB5 | 374786 | ENSG00000176927 |
| RNF148\|378925 | RNF148 | 378925 | ENSG00000235631 |
| TSPYL6\|388951 | TSPYL6 | 388951 | ENSG00000178021 |
| C2orf78\|388960 | C2orf78 | 388960 | ENSG00000187833 |
| PAGE2B\|389860 | PAGE2B | 389860 | ENSG00000238269 |
| BCAR4\|400500 | BCAR4 | 400500 | ENSG00000262117 |
| C1orf141\|400757 | C1orf141 | 400757 | ENSG00000203963 |
| C4orf40\|401137 | C4orf40 | 401137 | ENSG00000187533 |
| VCX3B\|425054 | VCX3B | 425054 | ENSG00000205642 |
| LRRC52\|440699 | LRRC52 | 440699 | ENSG00000162763 |
| CT45A4\|441520 | CT45A4 | 441520 | ENSG00000228836 |
| CT45A4\|441520.1 | CT45A6 | 441520 | ENSG00000226907 |
| CT45A4\|441520.2 | CT45A2 | 441520 | ENSG00000242185 |
| RFPL4B\|442247 | RFPL4B | 442247 | ENSG00000251258 |
| SPANXN5\|494197 | SPANXN5 | 494197 | ENSG00000204363 |
| CT45A1\|541466 | CT45A3 | 541466 | ENSG00000232417 |
| CT45A1\|541466.1 | CT45A1 | 541466 | ENSG00000232478 |
| RAD21L1\|642636 | RAD21L1 | 642636 | ENSG00000244588 |
| RHOXF2B\|727940 | RHOXF2B | 727940 | ENSG00000203989 |
| CT45A2\|728911 | CT45A4 | 728911 | ENSG00000228836 |
| CT45A2\|728911.1 | CT45A2 | 728911 | ENSG00000242185 |
| CT45A2\|728911.2 | CT45A1 | 728911 | ENSG00000232478 |
| CT45A2\|728911.3 | CT45A3 | 728911 | ENSG00000232417 |
| GAGE8\|100101629 | GAGE2E | 100101629 | ENSG00000205775 |
| GAGE8\|100101629.1 | GAGE2D | 100101629 | ENSG00000240257 |
| SPANXB2\|100133171 | SPANXB1 | 100133171 | ENSG00000235604 |
| SPANXB2\|100133171.1 | SPANXB2 | 100133171 | ENSG00000227234 |

**TS2:**
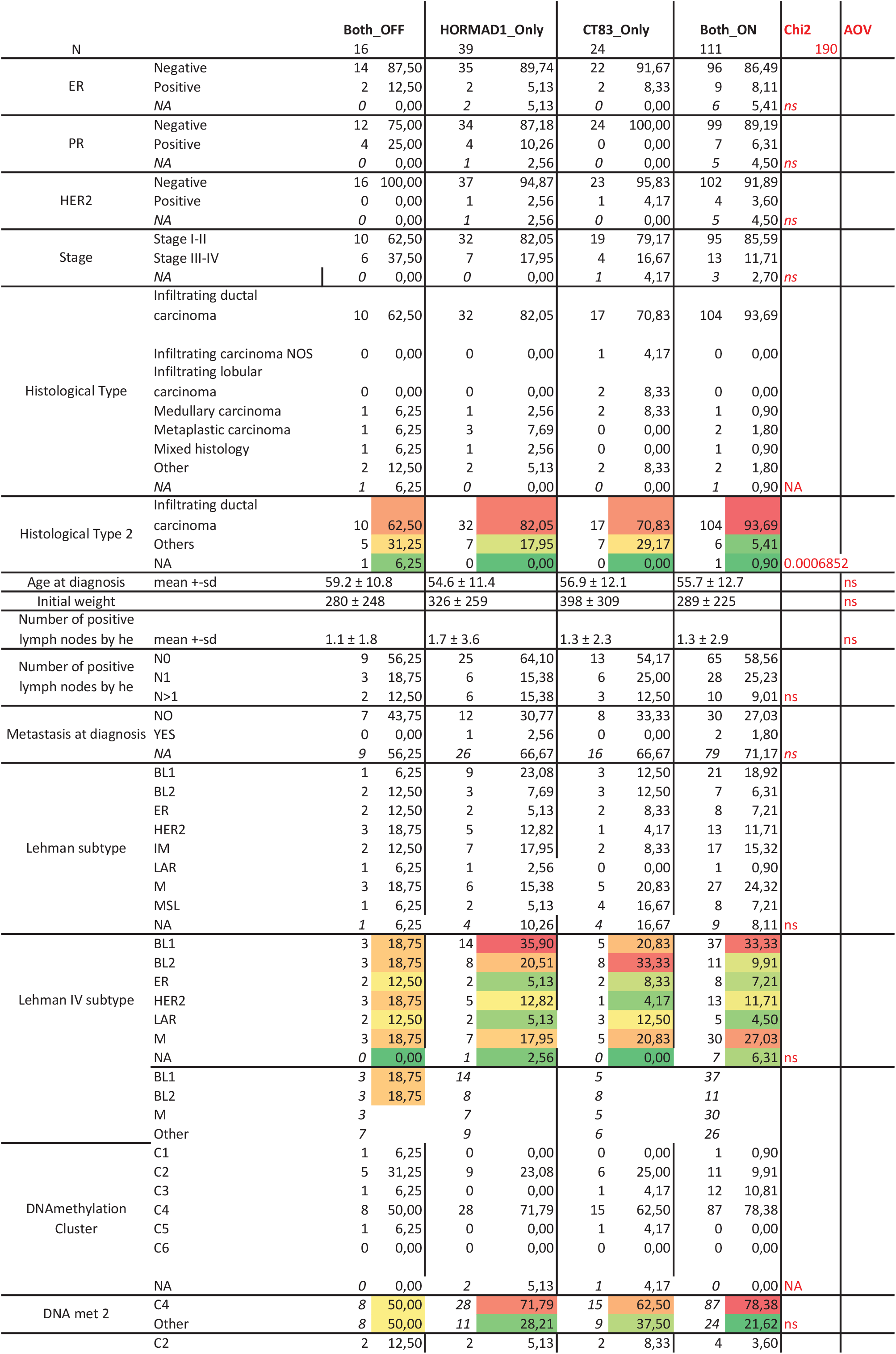
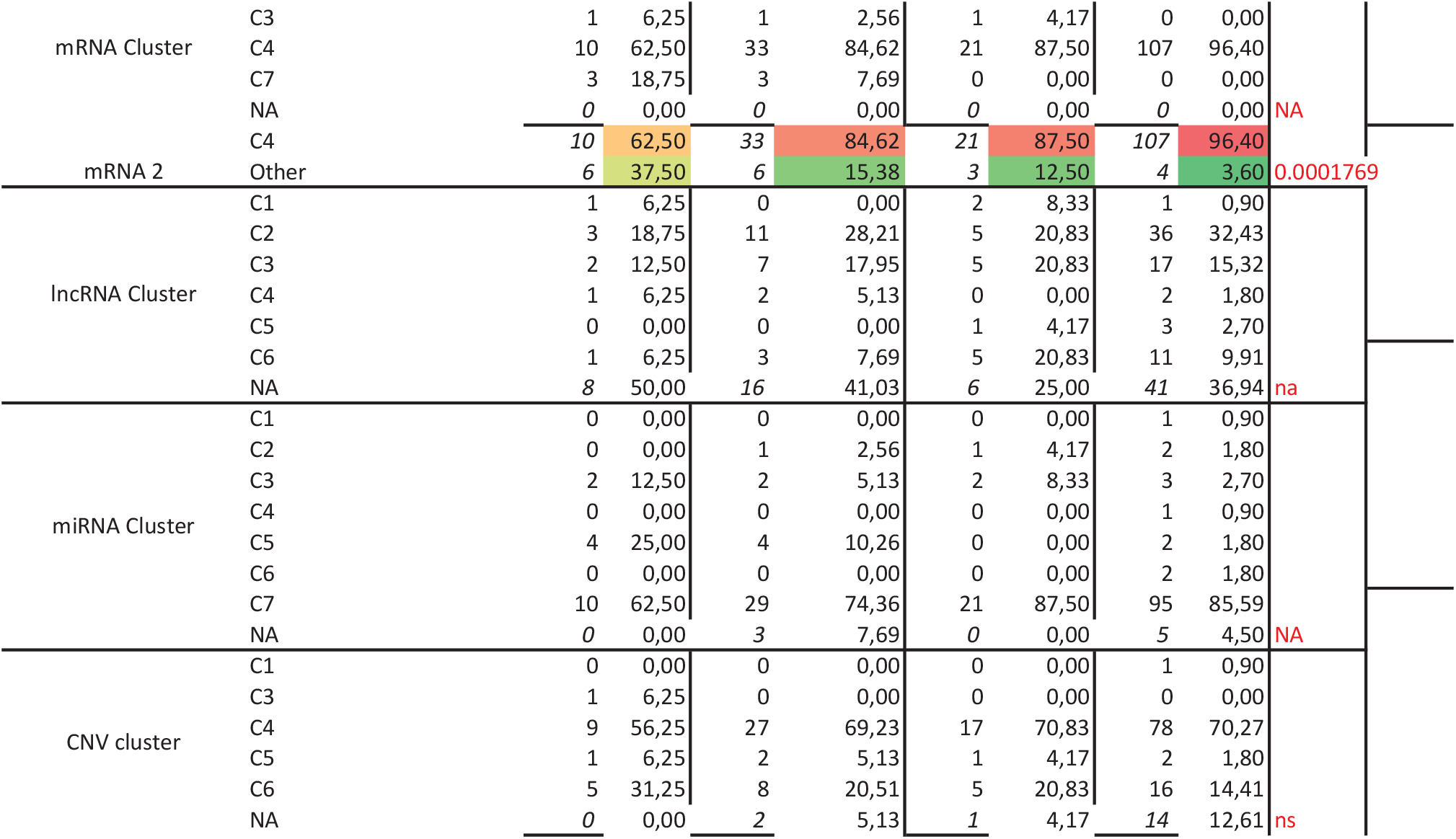
Basal-like tumor characteristics

**TS3:**
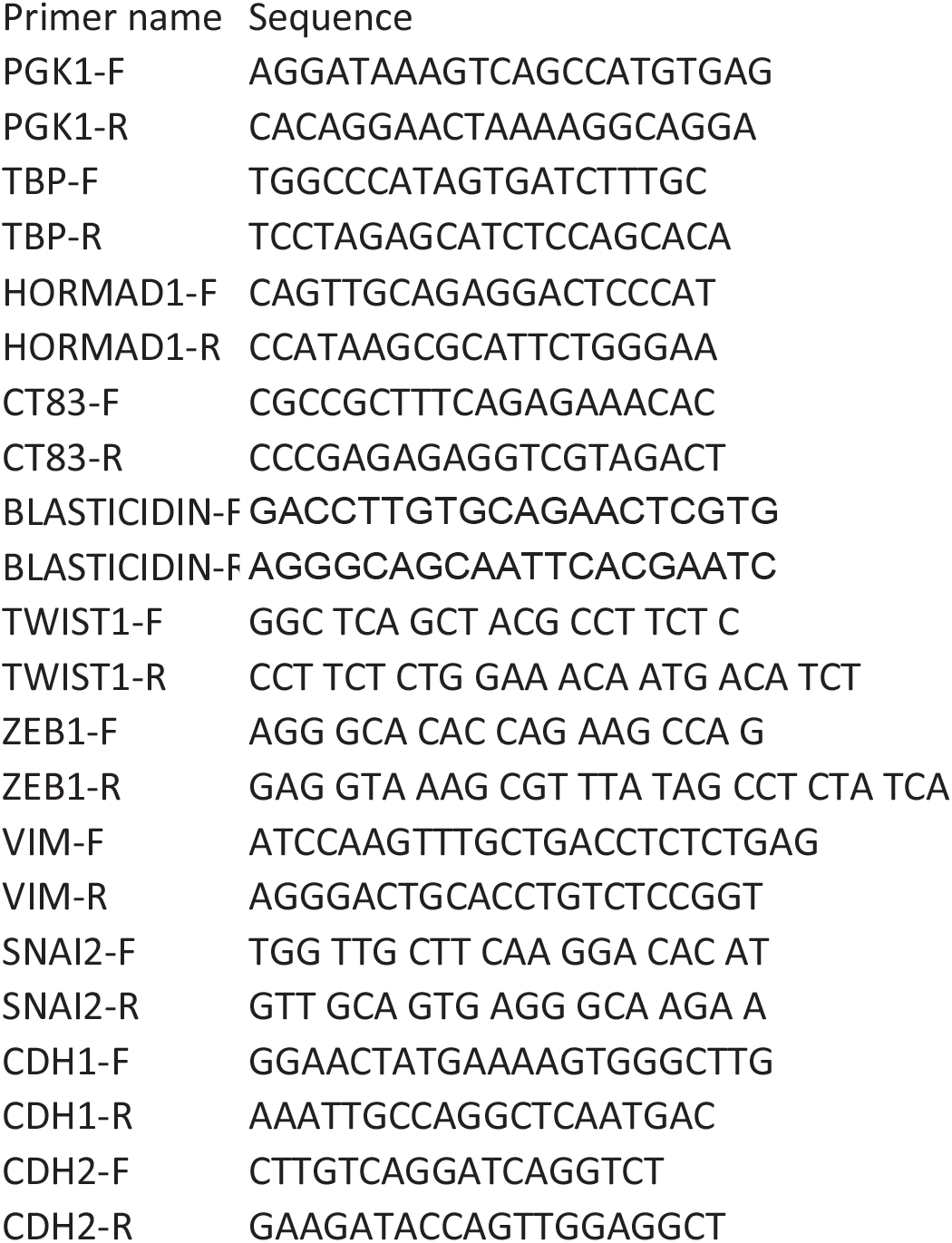
Primers

**TS4:**
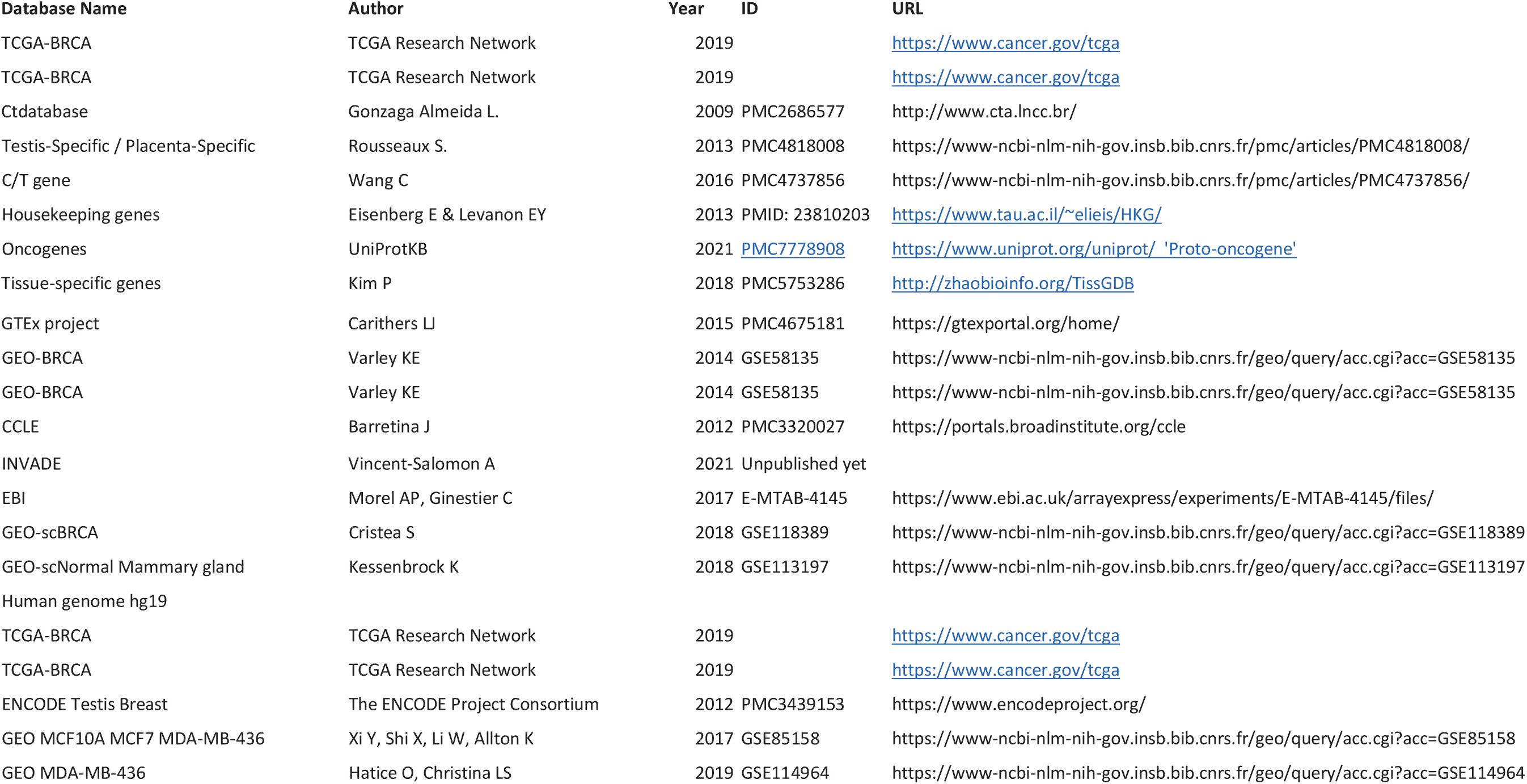
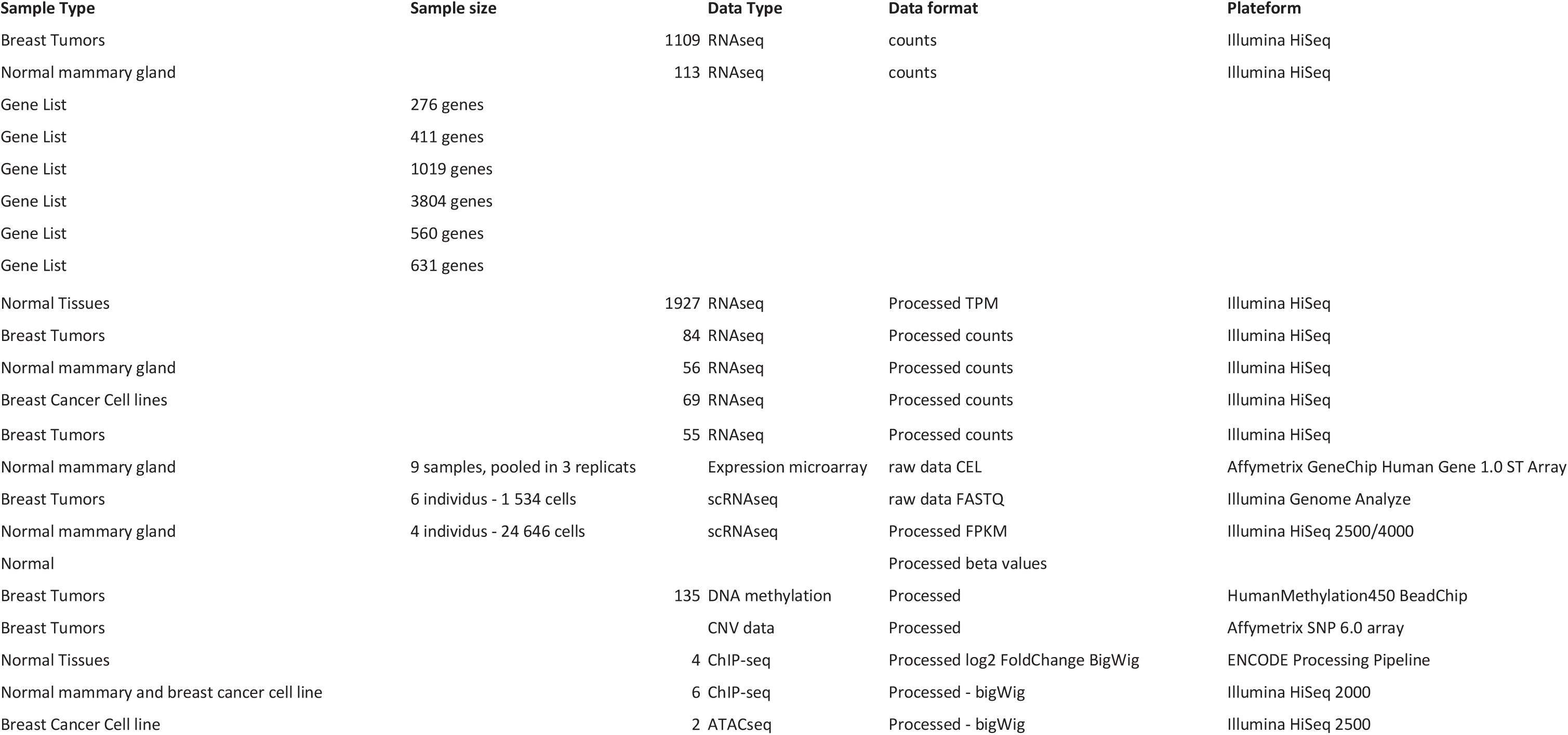
Dataset

